# Dynamic Instability of Dendrite Tips Generates the Highly Branched Morphologies of Sensory Neurons

**DOI:** 10.1101/2021.10.13.464245

**Authors:** Sonal Shree, Sabyasachi Sutradhar, Olivier Trottier, Yuhai Tu, Xin Liang, Jonathon Howard

**Affiliations:** Department of Molecular Biophysics & Biochemistry, Yale University, New Haven, CT 06511; Department of Physics, Yale University, New Haven, CT 06511; IBM T.J. Watson Research Center, Yorktown Heights, NY 10598; Tsinghua-Peking Joint Center for Life Sciences, School of Life Sciences, Tsinghua University, 100084 Beijing, China; Quantitative Biology Institute, Yale University, New Haven, CT 06511

**Keywords:** Branching morphogenesis, neuronal development, dendrite, arbor, dynamic instability, agent-based model

## Abstract

The highly ramified arbors of neuronal dendrites provide the substrate for the high connectivity and computational power of the brain. Altered dendritic morphology is associated with neuronal diseases. Many molecules have been shown to play crucial roles in shaping and maintaining dendrite morphology. Yet, the underlying principles by which molecular interactions generate branched morphologies are not understood. To elucidate these principles, we visualized the growth of dendrites throughout larval development of *Drosophila* sensory neurons and discovered that the tips of dendrites undergo dynamic instability, transitioning rapidly and stochastically between growing, shrinking, and paused states. By incorporating these measured dynamics into a novel, agent-based computational model, we showed that the complex and highly variable dendritic morphologies of these cells are a consequence of the stochastic dynamics of their dendrite tips. These principles may generalize to branching of other neuronal cell-types, as well as to branching at the subcellular and tissue levels.

## Introduction

Neurons are polarized cells (Cajal, 1906) whose axons and dendrites are often highly branched. Branching provides the surface area necessary for dendrites to receive inputs from thousands of other cells or from the environment (Lefebvre et al., 2013), and for axons to output signals to multiple cells. In these ways, branching facilitates the high connectivity of the brain (Lefebvre et al., 2013). Thus, the morphology of the neurons, together with their synaptic connections (Fornito et al., 2015; Zheng et al., 2018), defines the structure of the nervous system, the connectome, which is viewed as a prerequisite for understanding brain function (Denk et al., 2012). Whereas much is known about the growth of axons, whose tips, the growth cones, are guided by extracellular signals and guidepost cells (Stoeckli, 2018), the mechanisms underlying the growth and branching of dendrites are poorly understood. Elucidation of these mechanisms is the goal of the present work.

While many molecules have been shown to play crucial roles in shaping dendrites, the underlying rules by which molecular interactions generate branched morphologies is not understood. To investigate these rules, we have focused on dendrite morphogenesis in class IV dendritic arborization (da) neurons in *Drosophila,* a model system for dendritogenesis (Jan and Jan, 2010; Singhania and Grueber, 2014). These nociceptive neurons form a highly branched meshwork just under the cuticle that senses puncture of the larva by the ovipositor barbs of parasitic wasps and initiates avoidance behaviors (Basak et al., 2021; Robertson et al., 2013). Class IV cells are ideal for studying branching morphogenesis because they grow rapidly over 5 days of larval development, their branches are non-crossing due to self-avoidance mediated by the Down’s syndrome cell adhesion molecule (Hughes et al., 2007; Matthews et al., 2007; Soba et al., 2007) and other molecules (Emoto et al., 2004; Parrish et al., 2009), and they can be visualized using cell-specific labeling (Grueber et al., 2003; Jan and Jan, 2010). Many molecules that participate in dendrite morphogenesis have been identified: transcription factors (Jinushi-Nakao et al., 2007); extracellular matrix and integrins (Han et al., 2012; Kim et al., 2012); actin-associated proteins (Stürner et al., 2019); microtubule motors such as dynein and kinesin (Satoh et al., 2008; Zheng et al., 2008); microtubule regulators such as spastin (Sherwood et al., 2004), katanin (Stewart et al., 2012) and *γ*-tubulin (Ori-McKenney et al., 2012); and microRNAs such as *bantam* (Parrish et al., 2009). A major difficulty, however, is that it is currently not possible to predict quantitatively how developmental processes occurring at the molecular and subcellular levels determine the morphology of the entire dendritic arbor.

While several theoretical and computational models can produce dendrite-like branched morphologies, they are not grounded in molecular or development data. Early models, designed to describe and classify neurons, reconstituted morphologies based on the statistical properties of the observed arbors themselves (Ascoli and Krichmar, 2000; Nanda et al., 2018). Optimization-based models that minimize wiring (i.e., the total lengths of the branches) capture key features of neuronal morphology (Baltruschat et al., 2020; Cuntz et al., 2010), but lack connection to the cellular processes, as do models based on more abstract processes such as diffusion-limited aggregation (Luczak, 2006) and Turing-like pattern formation (Sugimura et al., 2007). More realistic models of *Drosophila* sensory cells, for example, capture important properties of the dendrite morphologies but use hypothetical branching and growth parameters (Ganguly et al., 2016; Palavalli et al., 2021). Models of branching morphogenesis in tissues are of limited applicability to dendrites: models of branching in the lungs (Metzger et al., 2008) and kidneys (Lefevre et al., 2017; Short et al., 2014) produce stereotyped morphologies that are distinct from the highly variable morphologies of neurons (Kanari et al., 2018). Stochastic models developed for other tissues, such as the mammary glands, use properties that are specific to these systems, such as tip bifurcation (Hannezo et al., 2017). Thus, current computational models fall short in providing a mechanistic understanding of dendrite morphology.

To circumvent these limitations, we have formulated a computational model that is based entirely on experimentally observed properties of dendrites measured over their development. The data-based model takes as input tip-growth dynamics, branching rates and self-avoidance measured using high-resolution, live-cell imaging in the developing animal. The model successfully recapitulates class IV dendrite morphogenesis and shows how the complex and variable morphology of dendritic arbors emerge from the microscopic dynamics of dendrite tips and provides insights into several mutant phenotypes.

## Results

As *Drosophila* larvae grow (Figure 1A), the arbors of their class IV dendrites also grow (Figure 1B). By the end of larval development, the meshwork of branches covers the larval surface like chainmail and the individual dendritic arbors fill the eight abdominal segments with widths up to 500 μm. Each abdominal segment on the dorsal side, on which we focus, has two class IV neurons, one on either side of the dorsal midline (see Figure 1C and Figure S1A for definitions of the larval axes), and each occupying an approximately rectangular hemisegment. When the first instar larva hatches at 24 hr after egg-lay (egg-lay is defined as time zero), the widths of the dendritic arbors (green and blue points) are smaller than the hemisegments (solid lines) as shown in Figure 1D (see Methods for how the sizes of the arbors and segments were calculated). Over the next 24 hr, the arbors grow faster than the segments and reach the edges of the adjacent hemisegments. By 72 hr, the arbor has densely filled the hemisegment (Figure S1B) and thereafter grows with the growing hemisegment in a process called scaling (Parrish et al., 2009). The tiling of the larval surface (Grueber et al., 2002), during which the dendrites do not cross into the adjacent hemisegments, is due to inhibitory interactions between neighboring class IV cells (Soba et al. 2007) and interactions with the adjacent epithelial cells (Parrish et al., 2009). We sought to understand how the dendrites grow and fill the hemisegments.

**Figure 1:**
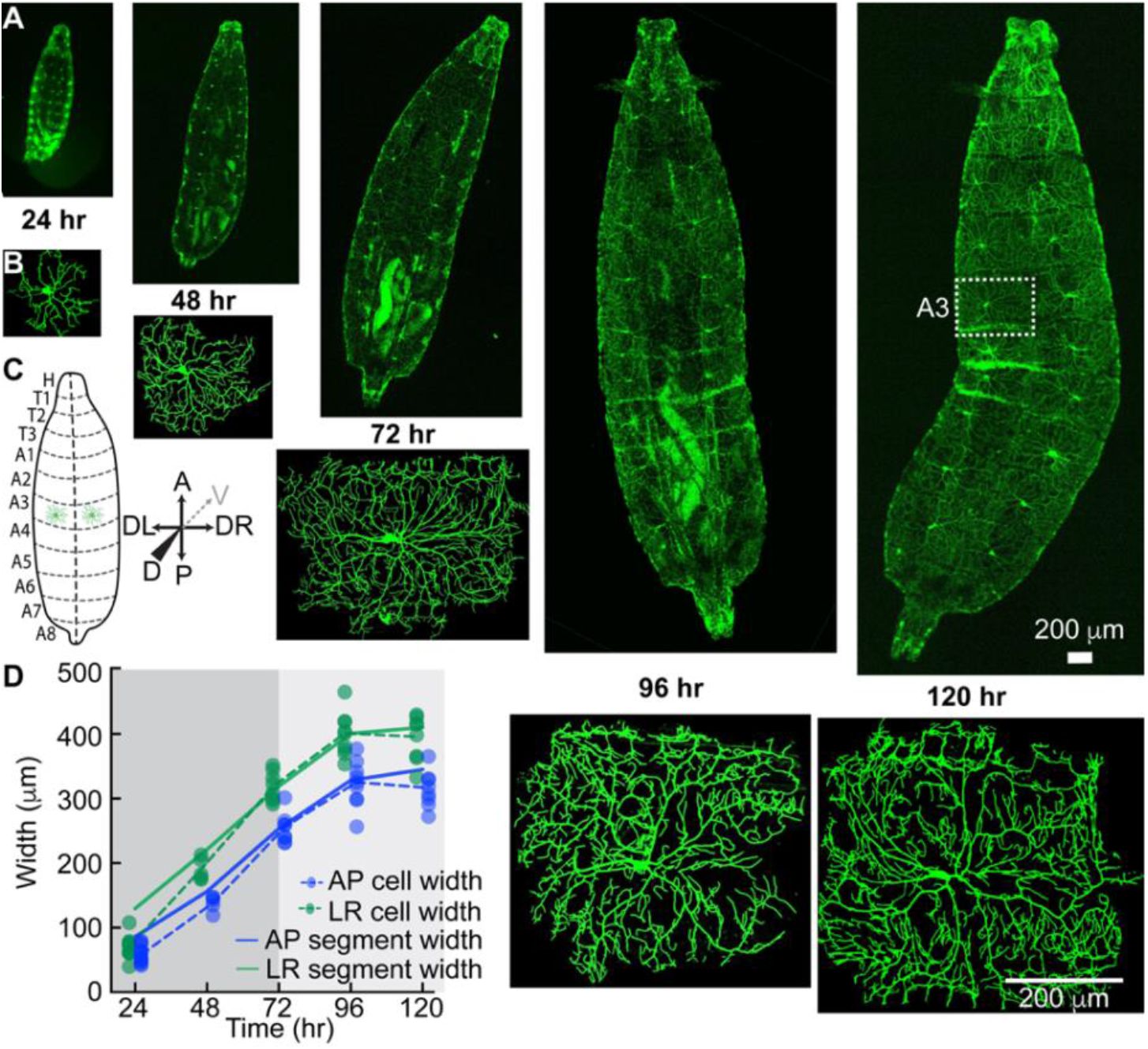
Growth of larvae and class IV neurons over development. **A** Whole-mount, living larvae imaged by spinning-disk confocal microscopy at 24 hr to 120 hr (egg-lay defined as time zero). Class IV neurons are marked with the transmembrane protein CD4 tagged with GFP (genotype - ;;ppkCD4-tdGFP). **B** Individual class IV cells from the A3 or A4 segments. An A3 segment is outlined in A (120 hr). **C** Cartoon of larvae as viewed from the dorsal side. The dashed line is dorsal midline. Anterior (A) is up and posterior (P) down. Left (DL) and right (DR) as viewed from the dorsal side (for the sake of simplicity we will mention DL-DR as LR everywhere in text and subsequent figures); the gray dashed arrow points in the ventral direction. **D** Growth of class IV arbors compared to their hemisegments. At 24 hr, the cell widths (solid circles with dashed lines through the averages) are smaller than the hemisegment widths (solid lines). In the next 24 hr, they touch the growing segment boundaries and by 72 hr (gray) they fill the hemisegment and then continue to grow with the hemisegment. The cell widths along each axis are defined as the sides of the rectangle which contains the same mass of branch skeleton distributed uniformly (see Methods).

### Dendrite growth is not due to elongation of all branches in the arbor

We first asked whether class IV arbors grow through the elongation of all their branches, both internal and terminal (Figure 2A, upper panel). In other words, does the arbor expand uniformly as shown in Figure 2A (middle panel), as proposed by (Yang and Chien, 2019). Such uniform expansion describes the growth of the overlying epithelial cells, whose number remains constant over larval development (Parrish et al., 2009), and of class I cells, which expand concomitantly with the segments (Castro et al., 2020; Palavalli et al., 2021). To test the role of branch elongation in arbor expansion, we reimaged the same neurons at discrete times over development, at 24 & 48 hr, and at 48 & 96 hr (Figure 2B). As the arbors grow, there is continuous addition and removal of branches. Nevertheless, it was possible to identify shared structural features of internal branches in the proximal region (Figure 2 C,D and Methods). The fractional increases in lengths of these identified internal branches were considerably less than the fractional increase in length of the hemisegments along the AP and LR axes (Figure 2E), in agreement with earlier measurements (Baltruschat et al., 2020). Because elongation of internal branches contributes only 3% (24-48 hr) to 11% (48-96 hr) of the overall growth of the dendrites, other mechanisms must contribute to the bulk of arbor growth. This finding implies that the proximal branches are not rigidly attached to the adjacent epithelium but must slowly slip as the hemisegments grow, an interesting issue that we will not explore further here.

**Figure 2:**
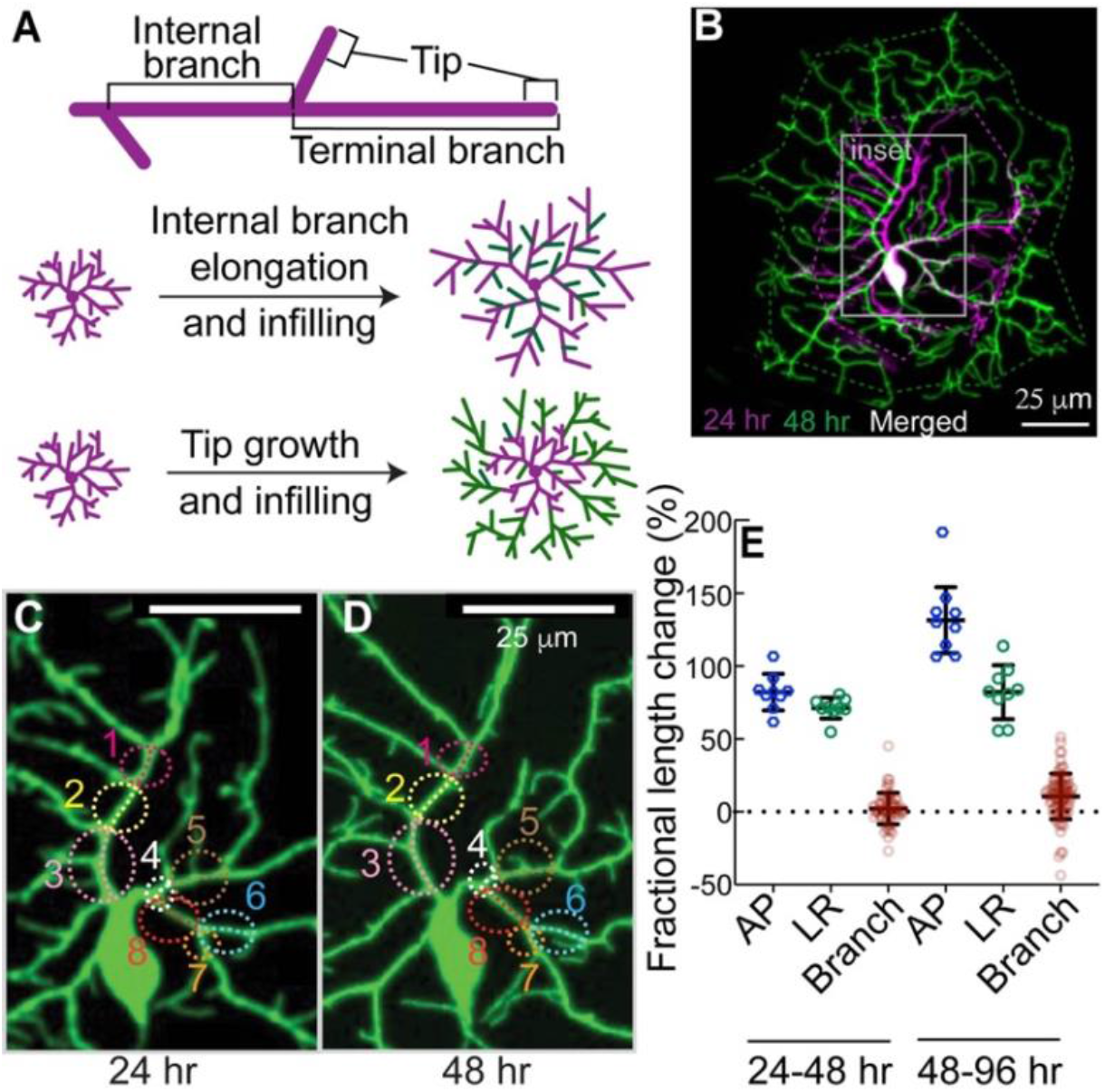
Branch dilation does not account for dendrite growth. **A** Upper panel: internal branches are branches that lie between two branch points as distinct from terminal branches that end in a tip. Lower two panels: two models for dendrite growth. Middle: elongation of existing branches & infilling with new branches. Lower: maintenance of internal branches & growth and infilling with new branches. **B** Maximum-projection image of a GFP-labeled class IV neuron cell (genotype - ;;ppkCD4-tdGFP) at 24 hr (magenta color) and same cell imaged at 48 hr (green color). 24 and 48hr images are combined with a leftward displacement of the latter. The area in the gray boundary (inset) is enlarged in **C** and **D** where conserved internal branches are marked with the same color circle and number (see Methods). **E** The fractional length changes of the hemisegments along the AP and LR axes were calculated from 24 hrs to 48 hrs and from 48 hrs to 96 hrs, together with the fractional length changes of the internal branches. Each blue and green circle is a different larva; the red circles correspond to several branch measurements in each of 6 larvae.

### Terminal dendrites grow from their tips and not from their bases

An alternative hypothesis to elongation of internal branches is that the growth of the arbor is due to branching and subsequent lengthening of the newly formed terminal branches (Figure 2A, lower panel). To test this hypothesis, the behavior of terminal dendrites was examined. Following birth by lateral branching from existing branches (Gao et al., 1999), terminal dendrites can: lengthen; shorten spontaneously or following contact with another branch; and pause (Figure 3A). Time-lapse imaging (Movies S1-S5) suggests that lengthening is due primarily to the addition of material near the tip. For example, the distances between the base of a branch and new branch points or bends do not change while the distal tips grow and shorten (Figure 3A, top-left panel; Movies S6-10). These observations argue against growth at the base and against uniform elongation along the length of terminal dendrites. Because new branches can form as close as 2 μm to an existing tip, we estimate that growth occurs within ~2 μm of the distal ends. Thus, tip growth, which occurs on timescales of minutes, may contribute to the overall growth of the arbor.

**Figure 3:**
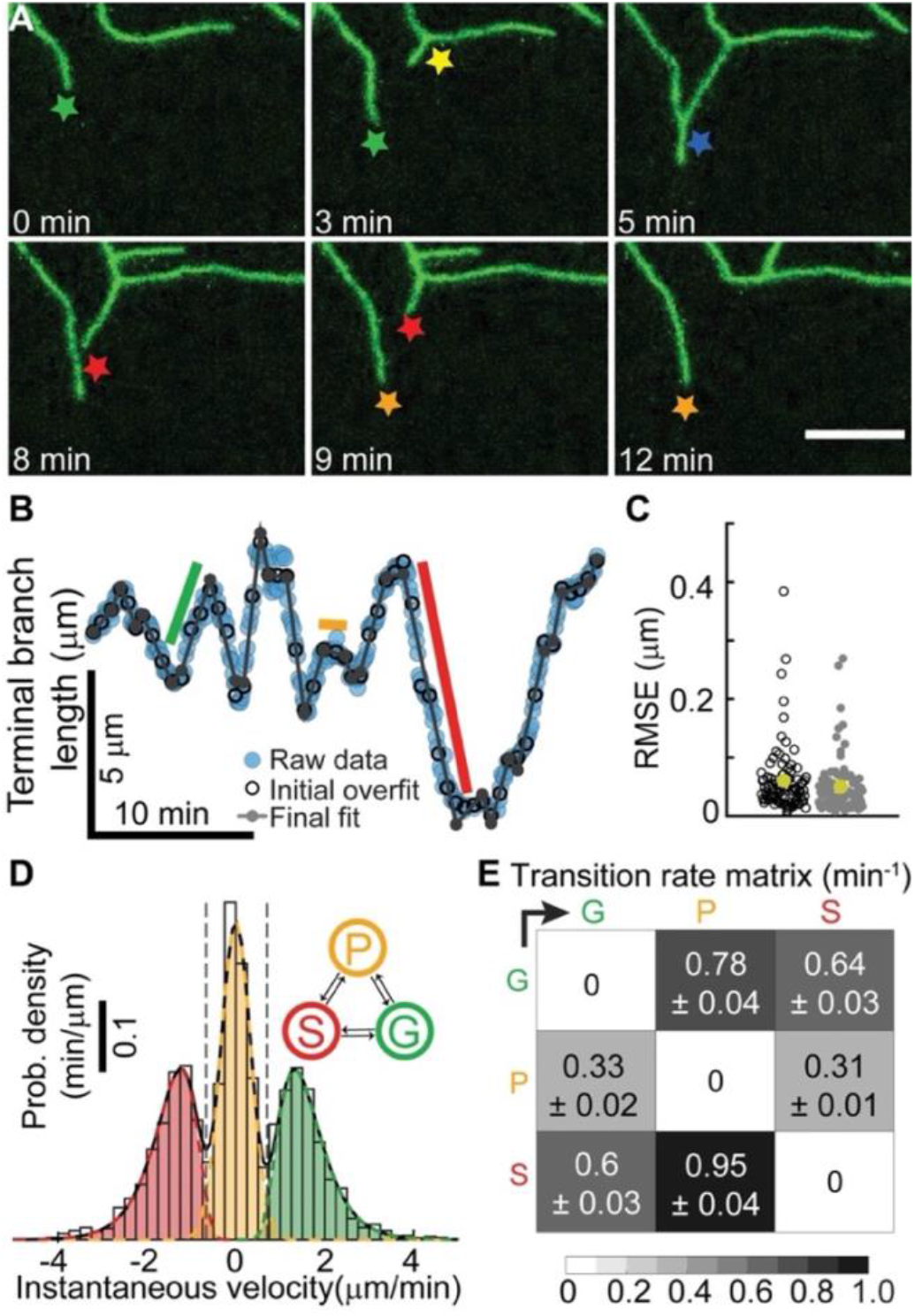
Dendrite tips transition between growing, shrinking and paused states. **A** Class IV dendrites growing (green star), birthing (yellow), colliding (blue), shrinking (red), and pausing (orange). Maximum projection spinning disk images of neurons were collected every 5 seconds for larval stage 24hr (genotype - ;;ppkCD4-tdGFP). **B** The length of a dendrite as a function of time (see Methods). The black open circles represent the initial piecewise linear fit using *N*/6 segments, where *N* is the total number of frames. The gray dots show the fitting after the iterative merging process (see Methods). Green, orange, and red indicated periods of growth, pausing, and shrinkage. **C** The root-mean-squared error before (black) and after (gray) merging for 91 trajectories (24 hr). The average error is 50 nm. **D** The velocity distribution shows three distinct peaks representing the growing (G), paused (P), and shrinking (S) states. **E** Transition rates between the three states.

### High-resolution tracking shows that tips transition between periods of constant growth velocity

To determine whether tip growth can account for arbor expansion, we tracked the lengths of terminal dendrites over time with an accuracy of ~0.1 μm (Methods, Figure S2 A-F). To ensure that mounting and imaging larvae did not interfere with growth, we restricted imaging to 20- to 30-minutes (Figure S1C). Typical trajectories show that dendrite growth is highly dynamic, with large fluctuations in velocity (Figure 3B, and Figure S2 G, I, K). To analyze tip trajectories, we first considered tip growth as a diffusion-with-drift process, a common way to describe particles moving in a flow. However, we ruled out this description because there were extended times of near-constant velocity: the green, red, and orange lines in Figure 3B (and Figure S2 G, I, K) clearly indicate periods of elongation, shortening, and stationarity. We therefore fit the trajectories to a piecewise linear continuous function, for which fast algorithms exist (D’Errico, 2021). This initial segmentation into regions of constant velocity provided a good fit to the tracking data (Figure 3B,C, black circles), showing that tip trajectories can be decomposed into sequential periods of linear growth or shrinkage.

### Dendrite tips undergo dynamic instability

We then asked the more difficult question of whether tips undergo dynamic transitions between growing (G), shrinking (S), and paused (P) states. In other words, can we classify the regions of constant growth into just three states such that transitions only occur between different states. Such a description is analogous to the dynamic instability of microtubules (Mitchison and Kirschner, 1984), which transition stochastically between two states: growing and shrinking.

To test whether a three-state dynamic model could account for tip growth, we assigned each region to be in a growing, shrinking or paused state by fitting the histogram of velocities with a three-peaked distribution, such as shown in Figure 3D, to define velocity thresholds between growth and pause, and shrinkage and pause. We then merged adjacent regions that belonged to the same state. Through an iterative procedure (see Methods), we segmented the trajectories into growing, shrinking, and paused states, with transitions only between different states. The resulting trajectory (Figure 3B, gray lines) was an excellent fit to the data: the root-mean-squared error was on average ~0.05 μm (Figure 3C), accounting for 85% to 99% of the variance (Figure S2 H, J, L). From these data, we calculated (i) the growing and shrinking speeds (Figure 3D) and (ii) the rates of the transitions between the three states (Figure 3E). At 24 hr, the growing and shrinking speeds were 1.61 μm/min and 1.52 μm/min, respectively, and the transition rates ranged from 0.31 to 0.95 per minute, corresponding to average lifetimes of individual states between 0.6 to 1.5 minutes. The net speed of dendrite elongation, ~0.034 μm/min (Table 1A), is much smaller than the average speed in the growing state because the dendrites spend roughly equal times in the growing and shrinking states.

**Table 1.** Dynamical parameters of dendrite tips at different developmental stages.

| **A: Pre-contact tip-dynamics parameters** |  |  |  |  |  |  |  |  |  |  |  |  |
| --- | --- | --- | --- | --- | --- | --- | --- | --- | --- | --- | --- | --- |
| Age (hr) | Tip velocity parameters |  |  | Tip transition rates |  |  |  |  |  | Corr. | Average tip velocity |  |
| | $\mu_G, \sigma_G$<br>($\mu\text{m}/\text{min}$) | $\mu_P, \sigma_P$<br>($\mu\text{m}/\text{min}$) | $\mu_S, \sigma_S$<br>($\mu\text{m}/\text{min}$) | $k_{GP}$<br>($\text{min}^{-1}$) | $k_{GS}$<br>($\text{min}^{-1}$) | $k_{PG}$<br>($\text{min}^{-1}$) | $k_{PS}$<br>($\text{min}^{-1}$) | $k_{SG}$<br>($\text{min}^{-1}$) | $k_{SP}$<br>($\text{min}^{-1}$) | $r \ \& \ p$ | $v_d^{Tracks}$<br>($\mu\text{m}/\text{min}$) | $v_d^{TranMat}$<br>($\mu\text{m}/\text{min}$) |
| 18-20 (E) | 0.37, 0.34<br>$V_G = 1.53$ | 0, 0.36 | 0.43, 0.37<br>$V_S = 1.65$ | 0.696 | 0.509 | 0.423 | 0.296 | 0.669 | 0.71 | 0.017<br>0.09 | 0.097<br>$\pm 0.0158$ | 0.100<br>$\pm 0.0151$ |
| 24 (L1) | 0.41, 0.36<br>$V_G = 1.61$ | 0, 0.34 | 0.35, 0.37<br>$V_S = 1.52$ | 0.784 | 0.64 | 0.335 | 0.314 | 0.598 | 0.946 | 0.008<br>0.65 | 0.034<br>$\pm 0.0196$ | 0.038<br>$\pm 0.0191$ |
| 48 (L2) | 0.40, 0.39<br>$V_G = 1.61$ | 0, 0.25 | 0.0, 0.41<br>$V_S = 1.1$ | 0.933 | 0.435 | 0.155 | 0.235 | 0.282 | 1.251 | 0.045<br>0.24 | 0.020<br>$\pm 0.1545$ | 0.026<br>$\pm 0.0177$ |
| 96 (L3) | 0.36, 0.52<br>$V_G = 1.64$ | 0, 0.25 | 0.19, 0.44<br>$V_S = 1.33$ | 0.923 | 0.799 | 0.116 | 0.117 | 0.575 | 1.276 | 0.002<br>0.9 | 0.008<br>$\pm 0.0039$ | 0.022<br>$\pm 0.0177$ |
| **B: Post-contact tip-dynamics parameters** |  |  |  |  |  |  |  |  |  |  |  |  |
| Age (hr) | Tip velocity parameters |  |  | Tip transition rates |  |  |  |  |  | Corr | Average tip velocity |  |
| | $\mu_G^*, \sigma_G^*$<br>($\mu\text{m}/\text{min}$) | $\mu_P^*, \sigma_P^*$<br>($\mu\text{m}/\text{min}$) | $\mu_S^*, \sigma_S^*$<br>($\mu\text{m}/\text{min}$) | $k_{GP}^*$<br>($\text{min}^{-1}$) | $k_{GS}^*$<br>($\text{min}^{-1}$) | $k_{PG}^*$<br>($\text{min}^{-1}$) | $k_{PS}^*$<br>($\text{min}^{-1}$) | $k_{SG}^*$<br>($\text{min}^{-1}$) | $k_{SP}^*$<br>($\text{min}^{-1}$) | $r \ \& \ p$ | $v_d^{Tracks}$<br>($\mu\text{m}/\text{min}$) | $v_d^{TranMat}$<br>($\mu\text{m}/\text{min}$) |
| 18-20 (E) | 0.53, 0.49<br>$V_G^* = 1.9$ | 0, 0.28 | 0.53, 0.54<br>$V_S^* = 1.9$ | 0.635 | 0.992 | 0.263 | 0.401 | 0.469 | 0.593 | 0.07<br>0.12 | -0.42<br>$\pm 0.07$ | -0.34<br>$\pm 0.07$ |
| 48 (L2) | 0.40, 0.50<br>$V_G^* = 1.7$ | 0, 0.25 | 0, 0.38<br>$V_S^* = 1.1$ | 1.446 | 1.24 | 0.134 | 0.29 | 0.239 | 0.814 | 0.018<br>0.76 | -0.23<br>$\pm 0.05$ | -0.19<br>$\pm 0.04$ |

Errors are SE.

This analysis shows that the growth trajectories accord with a three-state kinetic scheme, which provides a succinct yet comprehensive description of tip dynamics. This scheme is a generalization of the Dogterom and Leibler model of microtubule dynamic instability (Dogterom and Leibler, 1993), with inclusion of a third, paused state, and the growing and shrinking states having distributions of speeds.

### Dendrite dynamics and branching rates change over development time

Throughout development, the growing and shrinking speeds were roughly unchanged (Table 1A). The main change over development was that the transition rates out of the paused state decreased two- to four-fold and the transition rates into the paused state increased by about 50%. As a result, the dendrites spend more time in the paused state: they become less dynamic.

Branching, which always occurs on the sides of existing branches, also slowed down over development. The branching rate per unit dendrite length decreased roughly ten-fold from 24 to 96 hr (Figure 4A, Table S1). This decrease is another manifestation of dendrites becoming less dynamic over time. The geometry of branching, however, remained constant over development: the mean angle of a new (daughter) branch was close to 90° at all developmental stages (Figure 4B, Table S1), and the spatial distribution of branching remained roughly uniform (Figure 4C and D). In summary, both growth and branching are highly stochastic throughout development, though the transition and branching rates slow down over time.

**Figure 4:**
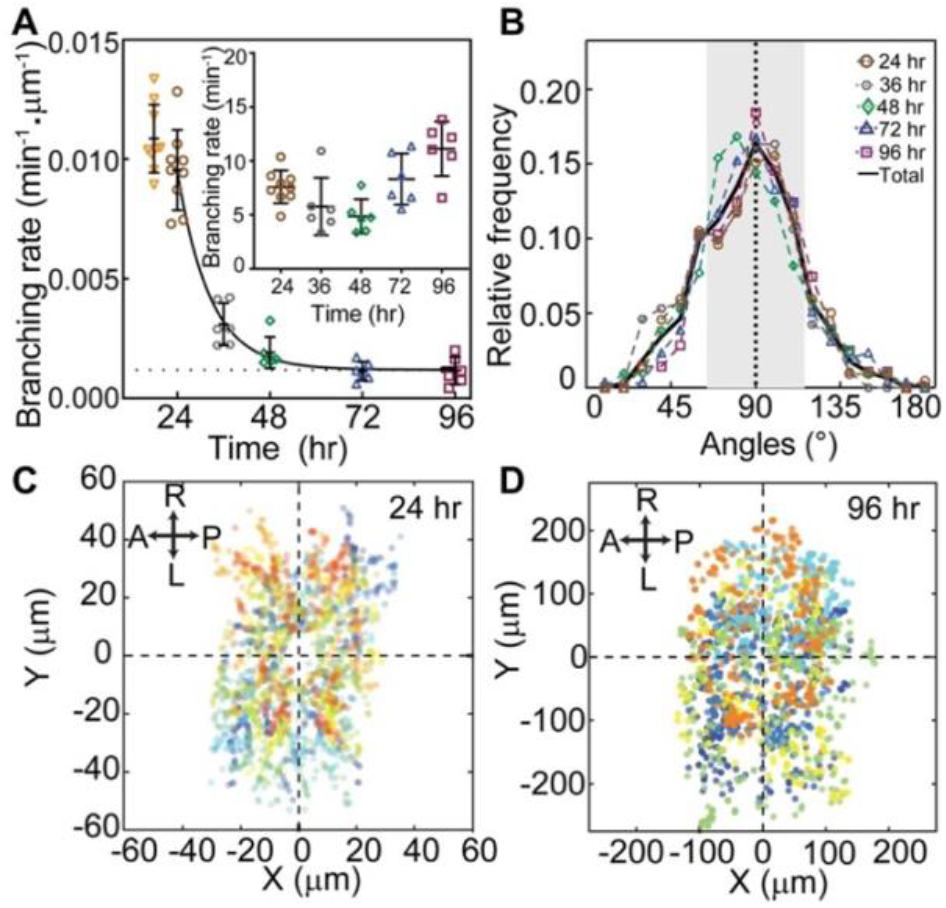
Dendrite branching over development. **A** The rate of appearance of new branches normalized by the total branch length is plotted against developmental time. Each symbol represents a neuron from a different larva (except at 24 hr where 3 neurons in each of 3 larvae were analyzed). The curve is an exponential fit with an offset (dotted line). Inset: The total branching rate per cell. **B** The distribution of branch angles between daughters at different developmental stages. The angle is zero when the new daughter grows parallel to the mother. Numbers of neurons: 6 (24 hr), 7 (36 hr), 4 (48 hr and 96 hr), 5 (72 hr) **C** Spatial distribution of branching events at 24 hr (9 neurons from 3 larvae). **D** Spatial distribution of branching events in 96 hr (6 neurons from 6 larvae). In both **C** and **D**, the soma positions are centered at the origin. Different colors represent different cells.

### Dendrite tips retain memories: their dynamics are not Markovian

We tested whether the transitions between growth states are Markovian, meaning that they depend only on their current state: i.e., they do not depend on the history, and there is no memory of earlier states. Consistent with a Markov process, the lifetimes of the states were approximately exponential at long times (Figure S3 A-I), and the probability of a transition in the sequence of occupied states (e.g., GPGSPGPSG...) did not depend on the previous state. For example, we found that the likelihood of G→P did not depend on the prior state: that is SG→P and PG→P were equally likely.

There were several violations of the Markov property, however. First, following contact of a tip with another branch, the growth dynamics change: the rates out of the growing state increase, so the dendrites spend less time growing (Table 1B). Therefore, in addition to contact-induced retraction, there is a long-lasting alteration of the dynamics: the average tip-growth rate changes from positive to negative (Table 1), and the post-contact dendrites shrink on average. This alteration implies that there is a long-lasting memory of the collision. Second, we found that the lifetimes of newly born dendrites were longer than expected for a Markov process. Following birth into the growing state, the transition rates, which were measured for older dendrites (>5 minutes after birth), predict that there will be an initial linear decrease in surviving dendrites due to the chance that a growing dendrite stochastically switches into a shrinking state, which then shortens and disappears. Instead, we found that the survival curve was initially flat, consistent with an initial growing state of 0.3 minutes (Figure S4A). Such a survival curve is another violation of the Markov process and implies that a newborn dendrite retains a memory of birth. Third, growth and shrinkage events with higher absolute speeds tended to have shorter lifetimes (Figure S3 J, K). Thus, while growth is highly stochastic, it deviates from being Markovian, indicating memory and “hidden variables” that influence the dynamics. These hidden variables are likely to be long-lived biochemical states (e.g., phosphorylation) triggered by dendrite birth or tip contact with other dendrites, which influence the dynamics.

### An agent-based model incorporates measured tip properties

To test whether the dynamical properties of the dendrites measured above can account for the observed morphology, we developed an agent-based computational model to predict morphologies based on tip dynamics. The elementary “particle” in the model is the dendrite tip, which serves as the agent (Figure 5A).

**Figure 5:**
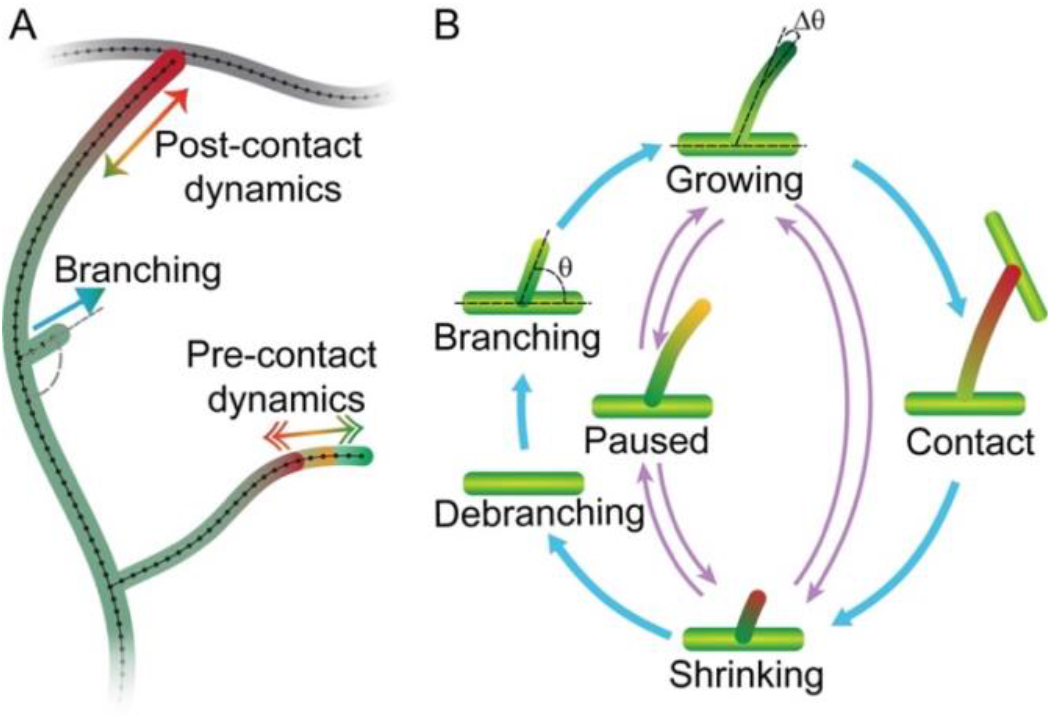
Schematic of the agent-based computational model. **A** Cartoon diagram depicted the different components of the model: tips are born by branching and transition between growing, shrinking, and paused states. Upon contact with another branch, the tips retract. **B** Diagram of the transitions. Parameters are listed in Table 1 and Tables S1-2.

Tips were simulated using rules that closely followed the experimental measurements (see Figure 5B, Methods). Terminal dendrites lengthen and shorten with speeds sampled from the growth and shrinkage distributions (e.g., Figure 3D). They transition between growing, shrinking, and paused states according to the measured transition rates (Table 1), which were linearly interpolated between different developmental stages. Tip birth occurred randomly in time and space along extant branches with the measured branching rates (Figure 4A). The nascent daughter branch was assumed to start in a growing state (Figure 5) with an initial length of 0.5 μm (Table S2) and included an initial lag of 0.3 min during which the transition out of the growing state was forbidden, in accordance with our observations. Dendrite death occurred when the last point disappeared during a shrinkage event. Contact, defined as a tip getting closer than 0.15 μm to another branch (roughly the radius of the terminal branch (Liao et al., 2021), switched a growing tip to a shrinking one with the post-contact dynamics (Table 1B).

The initial larval morphology at 24 hr was established by (i) growing two to four branches from a point (the origin) using the embryonic growth parameters at (18-20 hr) and (ii) allowing growth until the total branch length and number reached their values at 24 hr. To model the segment boundary, we assumed that contact with neighboring dendrites induced shrinkage (Parrish et al., 2009). Neighbor interaction was implemented using a periodic boundary condition such that one side of a growing neuron feels its opposite side as if growing on the topological equivalent of a torus. Over time, we gradually increased the size of the boundary according to the measured segment growth rates (Figure 1D).

### Simulated dendritic trees recapitulate coarse-grained features of dendrite morphology

We simulated dendritic trees using the parameters from Table 1 and Tables S1,2, all of which were measured or tightly constrained by experiments. The simulations (Figure 6A) recapitulated key properties of real arbors (Figure 6B).

i. **Arbor growth:** In the absence of a boundary, the widths of simulated arbors initially grew at 10 μm/hr, and then, after 48 to 72 hr, they slowed down to 4 μm/hr (Figure S5 Aii). Thus, the dendrite initially grew faster than the hemisegments (which grow at 4 μm/hour), leading to complete infilling by 72 hr; after 72 hr, arbor growth was just sufficient to keep up with segment growth. Contact-based retraction with the adjacent cell kept the dendrite confined to the hemisegment (i.e., tiling).
ii. **Total branch length and number**: The simulations predicted the observed increases in total branch length (Figure 6C) and number (Figure 6D), as well as the mean branch length (Figure 6E). The branch length distributions were roughly exponential in the simulations and the data (Figure S6 A-E). An interesting feature of the branch number is the initial burst (24-36 hr) and subsequent plateau (36-72 hr). The burst is predicted by the model and arises from two features of tip growth: the high initial branching rate (Figure 4A), and the perseverance of the initial growth of branches (i.e., a delay in transitioning out of the growing state). Without perseverance, which is a memory of birth, the plateau is less pronounced, showing that the initiation of branching is an important determinant of arbor morphology.

**Figure 6:**
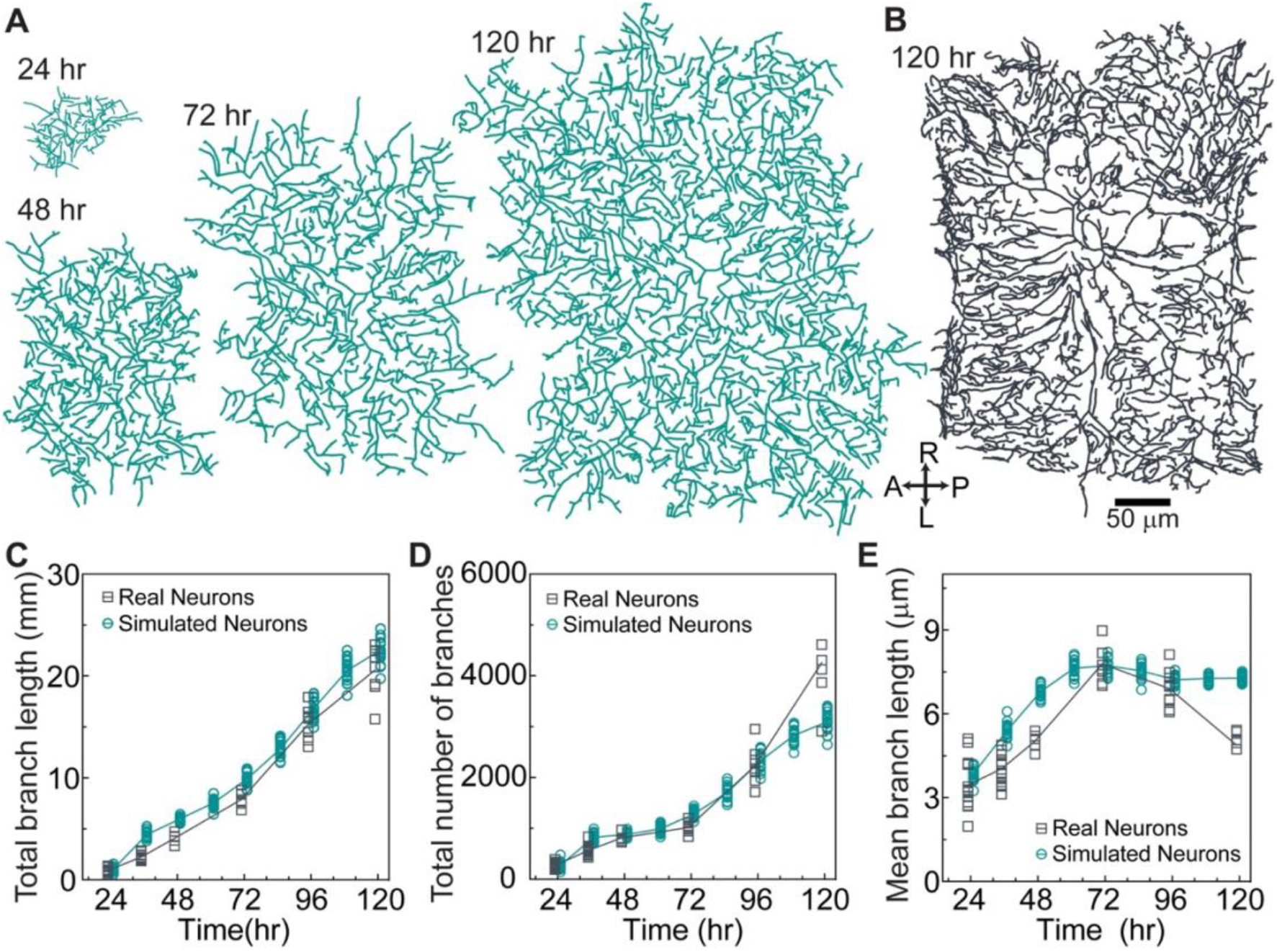
The agent-based model accords with overall neuronal growth. **A** Example of a simulated neuron using the parameters in Table 1 and Tables S1-2 at different developmental stages. Same scale bar as in B. **B** Example of the skeleton of a real neuron at 120 hr. **C** Total branch length over developmental for simulated and real neurons. D Total number of branches. **E** Mean branch length (total branch length/total number of branches).

There were some discrepancies between the data and the model. For example, the branch number of the simulated arbors saturated at 120 hr, while that of the real arbors continued to increase with an associated late decrease in mean branch length of real arbors. These discrepancies may indicate that dynamical properties change after 96 hr, the last time at which the dynamic parameters were measured (simulations beyond 96 hr are extrapolations). Another possible source of the discrepancy at 120 hours may be due the high branch density at intersegment boundaries along the AP axis (Figure 6B, right) arising from close cell-cell interactions, which were not predicted by the model. Nevertheless, we believe that the model is in good agreement with the average properties of the arbors.

### Simulated arbors capture the variability of dendrite morphology

In addition to predicting average branch properties, the model also predicted the variation of these features. For example, the measured branch numbers are highly variable, with coefficients of variation (SD/mean, CV) ranging from 0.11 to 0.22 over development. This CV is even larger than that of a Poisson process, a prototypical random process whose CV equals the inverse square root of the branch number (expected range 0.02 to 0.06). This comparison to a Poisson process indicates that branch number is highly variable from arbor to arbor, a manifestation of the stochasticity of the morphology. Total branch length and average branch length were also highly variable. Importantly, the simulations recapitulated this variability (Figure 6C-E). Thus, the model predicted both the average properties and the stochasticity of the branch number and length.

### Simulated arbors recapitulate fine-scale properties of dendrite morphology

The branches of both the simulated and real arbors formed dense meshworks (Figure 7A,B). We estimated the extent to which the branches cover the arbor using the box-counting method (Falconer, 1990) in which the number of boxes that contain a branch is plotted against the size of the boxes (Figure 7C). We found that the logarithm of box number was approximately proportional to the logarithm of box size, indicating that the patterns have scale-free and fractal-like properties. The proportionality breaks down at box sizes below 5 μm, the size of the “holes” in the pattern due to the average branch size. We defined the fractal dimension as the slope of the log-log plot (the power-law exponent) in the central region encompassing the middle 50% of the points (Figure 7C dashed lines). The fractal dimension increased from 1.4 to 1.8 from 24 hr to 120 hr for both the simulated and real arbors (Figure 7D). Because a region containing a single line has a fractal dimension of 1 whereas a region completely filled region has a fractal dimension of 2, the dendritic patterns are of intermediate dimension and at 120 hr nearly fill the plane (fractal dimension 1.8). Though the fractal dimensions of real arbors were consistently lower by about 0.1, we nevertheless conclude that the simulated arbors recapitulate the real arbors in this metric.

**Figure 7:**
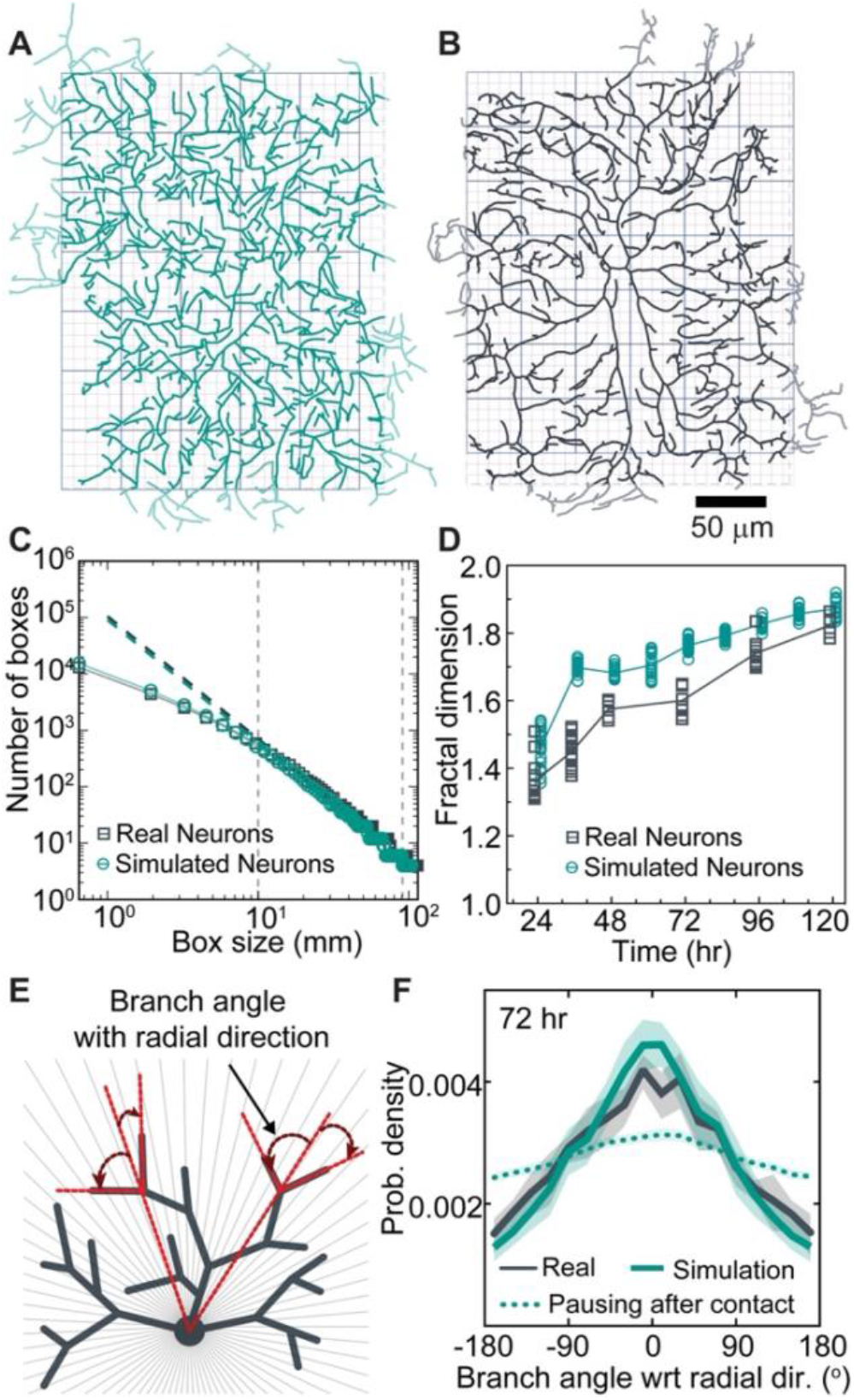
The agent-based model recapitulates the fine-scale patterns of real arbors. **A** Model arbor at 72 hr. Light green branches indicate the outer 5% of the arbors excluded from the analysis to mitigate against spurious boundary effects (see Methods). Boxes with two different sizes are shown in the background. **B** Skeletonized real arbor at 72 hr. Light gray branches were excluded from the analysis. **C** The number of boxes that contain a branch (*y*-axis) is plotted against the box size (*x*-axis). The slopes over the central 50% of the data (vertical dashed lines) define the fractal dimensions. **D** Fractal dimensions of real and simulated arbors both increase over developmental. **E** Diagram defining the radial orientation. **F** Radial orientation of a simulated tree (green solid line) and a real arbor (black) at 72 hr. Simulated tree with pausing instead of retraction (green dotted curve).

### Simulations recapitulate the radial orientation of dendrite branches

We discovered that class-IV cells have an unexpected long-range order: branches are not randomly oriented but instead tended to be parallel to the radial orientation (Figure 7E-F, Figure S6 F-J). The simulations also displayed radial orientation. Radial orientation is a consequence of contact-based retraction; if contact-based retraction is replaced by contact-based pausing, the radial orientation was greatly reduced (Figure 7F, dotted curve).

### Morphological predictions of the model

The agent-based model allowed us to explore which parameters are most important for arbor growth and mesh size (Figure S5) and provides hypothesis for the phenotypes of mutants.

A surprising finding was that branching drives overall arbor growth: increasing the branching rate not only increased the number and density of branches, as expected, but also increased arbor size (Figure S5B). Indeed, setting the average velocity of dendrite extension to zero still resulted in arbor growth, albeit slower (Figure S5E and Figure S7), showing that branching without net tip growth can drive arbor expansion. This is not to imply that the average tip growth rate is unimportant: doubling the net growth rate doubled the dendrite area (Figure S5D) and halving the net growth rate halved the dendrite area (Figure S5D). The latter finding accounts for the reduced arbors in Katanin (Kat-60L1) mutants, which spend less time in the paused states and more time in the shrinking state (Stewart et al., 2012): the mutant cells will therefore have a reduced net growth rate compared to controls, leading to smaller size (see Table S3).

Another surprising finding was that fluctuations in branch length also lead to growth. When the fluctuations were increased (by reducing the tip-transition rates) the growth rate increased, and vice versa (Figure S5C). This suggests an unexpected growth mechanism: length fluctuations are locked in by the formation of new branches, as only terminal branches can shorten and disappear. This stresses the importance of branching in growth.

The branching angle also affected arbor growth: if the branch angle was decreased to 45° (i.e., growth towards the direction of the mother branch), arbor growth increased, showing that outward growing branches are more likely to survive. The persistence of branch growth after birth was also important.

Our model shows dendrite density is set by the interplay between branching and self-avoidance. Branching is a form of positive feedback that increases branch density (Figure S5B). Therefore, branching is essential not just for expansion but for infilling the hemisegment. Self-avoidance is negative feedback: reducing self-avoidance in the model increases the branch density (Table S3), which is observed in studies in which self-avoidance molecules are mutated (Emoto et al., 2004; Hughes et al., 2007; Matthews et al., 2007; Soba et al., 2007).

By performing this variational analysis (e.g., Figure S5), we could identify which of the 67 parameters (see Methods) are key to overall dendrite growth and morphology. The key parameters are: the net growth velocity and its variance together with the net shrinkage following contact (3 parameters) and the branching rate (1 parameter) and angle (2 parameters). In the absence of the boundary, these determine the growth rate, the branch number and length, together with the fine structure (fractal dimension and radial orientation). While the detailed growth and morphology depend on the change of these 6 parameters over development (and the boundary), these parameters are the fundamental ones that specify growth and morphology.

## Discussion

We have discovered that the tips of *Drosophila* class IV dendrites transition stochastically between three states - growing, shrinking, and paused. This allows dendrites to explore extracellular space, analogous to the exploration of intracellular space by microtubules undergoing dynamic instability (Kirschner and Mitchison, 1986). Our modeling shows that these transitions, together with lateral branching, and contact-mediated retraction, give rise to the complex and highly variable morphology of the dendritic arbors, allowing them to fulfill their biological functions. The dense, almost unbroken meshwork optimizes detection of the fine ovipositor barbs of parasite wasps (Basak et al., 2021; Robertson et al., 2013). And the radial orientation of branches, a form of long-range order that emerges from local interactions (Toner and Tu, 1995; Vicsek et al., 1995), in this case contact-induced retraction, reduces the path distance to the cell body: this minimizes signaling delays and wiring costs (Baltruschat et al., 2020). Thus, stochastic tip dynamics may partially solve the riddle of how “the morphological features displayed by neurons appear to obey precise rules that are accompanied by useful consequences” (Cajal, 1995).

### Model limitations

The agent-based model fails to account for some features of class IV cells. For example, the model does not extrapolate well to 120 hr, suggesting that important developmental changes may occur after 96 hr. The model also fails to predict the asymmetry of real arbors, which form close contacts with class IV cells in the adjacent segment along the AP axis but not along the LR axis (Figure 6A,B); the model contains no asymmetries. Thus, the interactions between neighbors is more than just the contact-based retractions assumed in the model. Another shortcoming is that many simulated branches have sharp angles, whereas real branches are smoother: this is because when a mother branch shrinks back to a daughter in the model, the original branch angle is preserved (average 90 degrees), whereas in the real cells the bend smoothens over time (see an example of this in Supp. movie 7). Thus, there are important features of real class IV dendritic arbors that are not accounted for. Other important aspects of dendrite morphology, such as branch diameters (Liao et al. 2021) and the three-dimensionality of class IV cells (Han et al. 2012), have not been included in the model. Moreover, the model does not take into account the guidance of class IV dendrites by external cues such as the cuticular epithelium (Parrish et al., 2009; Uçar et al., 2021) and other neurons (Grueber et al., 2002), though theoretical tools have recently been developed to incorporate these cues (Parrish et al., 2009; Uçar et al., 2021). Finally, the internal branches in our model are completely immobile, whereas we have observed internal branch movements with respect to the substrate. Despite these limitations, however, our model provides a framework on which to build more complex interactions.

### Dendrite tips: an intermediate organizational principle of dendrite morphology

Our results strongly support the concept that the dendrite tip is a “branching engine … that initiates, directs, and maintains branch outgrowth during development and regrowth” (Lu and Werb, 2008). The dendrite tip, with a diameter of only 0.2 μm (Liao et al., 2021) and with dynamics on the timescale of ~1 minute (the state lifetimes), generates structures up to 500 μm in diameter (>1000-times larger sizes) over five days (>1000-times longer times). Tips, therefore, are intermediate in length- and timescales between molecules (small size and short-time scale motions) and morphology (large size and long-timescale motions).

The concept of the dendrite tip as a branching machine has four important implications. First, if the molecular basis of tip growth and branching can be elucidated, our agent-based model will provide a full connection between genotype and phenotype, with the caveats that morphogenesis is stochastic and some features such as three dimensionality are not included. Second, altered tip dynamics due to mutations and diseases may underly altered dendrite morphologies (see Introduction and the Predictions of the model section). Third, tip rules may specify neuronal identity, often defined by dendrite morphology (DeFelipe et al., 2013). And fourth, the stochastic nature of the tip rules may facilitate the evolution of neuronal cell types. This is because developmental stochasticity amplifies genetic variation by allowing a large class of morphologies to be sampled for each genotype. While some morphological outliers may function poorly, others might be beneficial, and genetic and/or epigenetic mechanisms could selectively stabilize these beneficial morphologies.

### Potential molecular mechanisms underlying dendrite tip dynamics and branching

Neurite elongation by tip growth also occurs in axons (Dent et al., 2011) and in other dendrites, such as those in *C. elegans* PVD neurons (Liang et al., 2020). An important difference between dendrite tips in class IV cells and the growth cones seen in these other cells is that the tips of class IV dendrites are much smaller. The diameter of class IV terminal dendrites is only ~200 nm (Liao et al., 2021), as small as a single filopodium, the finest feature of growth cones observed under the light microscope. Thus, tip-growth mechanisms in class IV dendrites likely differ from growth-cone-based growth. An important open question is how the cytoskeleton and membranes reach the dendrite tips: what are the relative contributions of diffusion, filament polymerization (Jinushi-Nakao et al., 2007; Ori-McKenney et al., 2012; Yalgin et al., 2015), motor-driven transport (Weiner et al., 2016) and motor-driven sliding (Winding et al., 2016)?

### General mechanisms of branching morphogenesis

Dendrite tips share features of branch tips in other systems. Tip cells drive branching in branched tissues such as the mammary glands (Lu and Werb, 2008). The growing ends of cytoskeletal filaments, with associated nucleation factors, drive branching of organelles such as the microtubule-based mitotic spindle (Decker et al., 2018; Petry et al., 2013) and the actin-based lamellipodium (Pollard and Borisy, 2003). In all three cases (dendrites, tissues and cytoskeleton), the tips operate at shorter length-scales and timescales than those of the structures they produce. Furthermore, they all respond to external signals: contact-based retraction of dendrite tips, cortex-induced catastrophe of microtubule ends (Komarova et al., 2002), and self-avoidance of mammary-gland branches (Lu and Werb, 2008). Finally, all three are stochastic and generate highly variable morphologies. Given these commonalities, it is likely that the principles that we have elucidated for dendrites generalize to other branched systems.

### Branching

Our observation that branching is an intensive property—the total rate of formation of branches is almost independent of arbor size (Figure 4A, inset)—is evidence that there are only a limited number of “branching factors” being produced in the entire cell per unit time. The uniform distribution of new branches suggests that the branching factors are dispersed throughout the cells, perhaps by molecular motors. Several phenotypes of molecular motor mutants in class IV cells support this hypothesis. The perturbation of molecular motors and their adapter proteins, including dynein (Arthur et al., 2015; Satoh et al., 2008; Zheng et al., 2008) and kinesins (Kelliher et al., 2018; Satoh et al., 2008), result in non-uniform branch densities, as expected if the distribution of branching factors were disrupted (see Table S3).

### Generalization of the agent-based model to other neurons

To explore the generality of our model, we have simulated different neuronal cell types, such as *Drosophila* class I neurons, mammalian retinal ganglion cells, Purkinje cells and starburst amacrine cells. By modifying the input parameters, our model was able to successfully capture the key morphological features of these cells as shown in the Figure S8. In class I cells, contact-based tip retraction leads to secondary branches being orthogonal to the primary branch even when the initial branching angles are uniformly distributed (Figure S8A); this confirms the finding of (Palavalli et al., 2021) and is related to the radial orientation of class IV cells described above. Contact-based retraction also leads to the radial orientation of retinal ganglion cells (Figure S8B), though we found better agreement using a small branching angle (45 degrees relative to the direction of the mother). To simulate Purkinje cells, we assumed slow tip growth of dendritic tips and complete retraction after contact to recapitulate the locally parallel branch orientations (Figure S8C). To simulate starburst amacrine cells, it was necessary to replace lateral branching with tip bifurcation (Figure S8D). Though our model can recapitulate certain aspects of the morphologies of these cells, these simulations are just predictions based on hypothetical model parameters and need to be tested experimentally. Nevertheless, these examples show that the model is versatile and has predictive potential beyond just *Drosophila* class IV sensory neurons.

If the dendrite branching rules deduced for class IV cells do indeed generalize to other neurons, then they may facilitate mapping connectomes by providing anatomical constraint on connectivity, as well as giving insight into genetic disorders that affect morphology (Forrest et al., 2018; Kapitein and Hoogenraad, 2015; Koleske, 2013; Kulkarni and Firestein, 2012).

## Materials and Methods

### Fly Stocks and maintenance

The fly line *;;ppk-cd4-tdGFP* (homozygous) was used to image class IV dendritic arborization neurons and was a kind gift from Dr. Chun Han (Cornell University). Fly crosses were maintained in fly chambers at 25 °C, 60% humidity in a Darwin Chamber with 12 -hour light/dark cycles. An apple-agar plate was used to collect the fly embryos and a big drop of yeast paste was put in the center of the agar plate to induce egg-laying.

Apple-agar plates were made by mixing 4X apple juice concentrate (355 ml), water (300 ml), dextrose (155 gm), sucrose (80 g). This solution was stirred and heated to dissolve the sugars, and agar (Bacto agar, Becton Dickinson, 60 g) and 1.25N NaOH (70 ml) were added. The solution was covered loosely with foil and autoclaved in the liquid cycle for 30 min. The plating mixture contained 100 ml of this apple-agar concentrate, 197 ml of water, and 3 ml of Acid mix A—an equal mixture of propionic acid (100%, 83.6 ml, and 16.4 ml water) and phosphoric acid (100%, 8.3 ml and 91.7 ml water).

For neuron morphometrics, embryos were collected every 15 minutes and imaged when they reached the appropriate age **A**fter **E**gg **L**ay (AEL): 24hr, 48hr, 72hr, 96hr, and 120hr.

### Sample preparation

For imaging, embryos of appropriate age (18 hr to 22 hr AEL) were collected from apple-agar plates and dechorionated by gently rolling them on a piece of double-sided tape stuck to a glass slide. The dechorionated embryos were then placed with their dorsal side down on a No. 1.5 coverslip, MatTex, with a small drop of halocarbon oil 700. A piece of wet Kim wipe was placed near the embryos to maintain humidity during imaging. No anesthetics were used for embryo imaging. For larvae imaging, larvae of ages 24 hr, 48 hr, 72 hr, 96 hr, and 120 hr were washed with 20% and 5% sucrose solution, anesthetized using FlyNap (Carolina Biologicals, Burlington, NC, USA), and transferred to apple-agar plates to recover for 1-5 minutes. After recovery, larvae were gently placed with their dorsal side up on a 1% agar bed adhered to a glass slide and imaged in a drop of 50% PBS, 50% Halocarbon oil 700 (Sigma Aldrich). Larvae were further immobilized by gently pressing them with a 22mm X 22mm coverslip lined with Vaseline or vacuum grease.

### Imaging

Samples were imaged on a spinning disk microscope: a Yokogawa CSU-W1 disk (pinhole size 50 μm) built on a fully automated Nikon TI inverted microscope with perfect focus, 488nm laser illumination at 18-21 % laser power, either a 40X (1.25 NA, 0.1615-micron pixel size) or a 60X (1.20 NA, 0.106-micron pixel size) water immersion objective, an sCMOS camera (Zyla 4.2 plus), and Nikon Elements software. The temperature of the sample region was maintained using an objective space heater at 25°C (OKO labs stage heater). Samples were manually focused to identify abdominal third and fourth segments (A3 or A4 neurons) before image acquisition. Full-frame movies (2048×2048 pixels) containing 6 to 12 1-μm sections were collected every 4-6 s. Static images for morphometric studies were acquired using a 60X water immersion objective for 24 hr and 40X objective for later stages. Images were stitched using in-house code (https://github.com/oliviertrottier/neuron-stitch). Movies were curated for subsequent offline analysis. Image analysis (segmentation, skeletonization, branching analysis, and angle measurements) was done using ImageJ.

### Controls for growth

Class IV neurons are susceptible to mechanical pressure, which, if too large, stops growth and causes degeneration. Therefore, embryos were imaged without an overlying coverslip. Larvae were immobilized with minimal pressure under a coverslip and were placed on a 1% agar bed. As a control, we plotted the size of neurons throughout imaging to confirm that they remain on the “standard” growth curve (Figure S1 C).

### Segment length determination

The *Drosophila* larval abdomen is divided into eight abdominal segments (Figure 1C & Figure S1A). Each segment on the dorsal side has two neurons, each occupying a hemi segment (Figure 1C). Each hemisegment is approximately rectangular with an Anterior-Posterior (AP) and Left-Right axis. The width of the segment along the AP axis was measured as the distance between the cell bodies of the adjacent neurons along the AP axis. The LR width was measured as the distance between cell bodies in adjacent hemisegments across the dorsal midline corrected for the offset of the cell bodies, which are not in the centers of the cells but displaced away from the midline. These segment widths were measured abdominal segments A2 to A5 for three larvae for all the respective stages.

### Arbor skeletonization and branch length measurements

Scanning-confocal images were maximally projected and individual dendrites manually segmented from their neighbors. The segmented neurons were binarized using a custom algorithm and skeletonized using MATLAB’s ‘`bwmorph`’. The individual branches were identified by subtracting the branch points identified by ‘`bwmorph`’ from the skeleton. The pixel coordinates of the branches were smoothed using a spline prior to calculating the total length of all branches and their average length.

### Dendrite arbor width

The arbors of class IV are approximately rectangular with axes parallel to the AP and LR axes. If the mass of the dendrite skeleton is uniformly distributed in a rectangle, then the widths, *D*_AP_ and *D*_LR_, are:

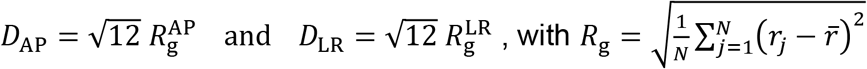

where *R_g_* is the radius of gyration, *N* is the total number of occupied pixels in the skeleton, *η* is the projection onto the respective axis of the *j*^th^ occupied pixel and 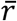 is the mean projected position of all occupied pixels. *R_g_* is the standard deviation of the dendrite pixels i.e., their spread from the center. We confirmed that the widths defined in this way were good approximations to the rectangles containing 95% of the skeletal mass.

### Analysis of the elongation of internal branches

To study the possible role that the elongation of internal branches in arbor growth, we imaged the same dorsal neurons (A3, A4, and A5) every 24 hrs. Larvae were mounted and imaged as described but without the use of anesthetics. Their movement was minimized by imaging at 4 °C for 2-5 mins. They were then returned to the apple-agar plate in the Darwin Chamber. The larvae were imaged using 20X and 40X objectives. For image analysis, the same neurons at 24 & 48 hr, and 48 & 96hr were segmented and aligned using ImageJ to identify conserved non-terminal internal branches in the proximal region. The fractional increases in branches and segment lengths were defined as

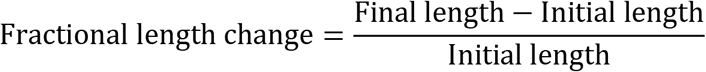

### Branching rate and branch angle

To determine branching events, time-lapse movies of duration 20-30 minutes were analyzed manually using ImageJ. A new protrusion of length >0.25 μm was scored as a new branch. The total branching rate (min^-1^) was calculated by dividing the total number of branching events by the total time. The specific branching rate (μm^-1^min^-1^) was calculated as the total branching rate divided by the total branch length. The spatial distribution of all branching events was plotted using MATLAB with the soma at the origin *(x = 0,y =* 0). The angle of new branches was measured using the angle tool of ImageJ (zero angle defined as in the direction of the mother). The angle distribution graph was plotted using Prism.

### Fractal Dimension

We used the box-counting method to calculate the fractal dimension. For each box width, *W,* we measured the number of boxes, *N*(*W*), needed to cover all the occupied pixels of the skeleton (Figure 7A,B). *N*(*W*) is approximately linear on a log-log plot (Figure 7C) indicative of a power law. The middle 50% of points (between the dashed lines in Figure 7C) was fit to *N*(*W*) = *W*^-D_f_^ to obtain the fractal dimension *D*_f_.

### Tip-tracking algorithm

To quantify the dynamical properties of the tips, we developed an in-house algorithm to track dendritic tips and determine dendrite length-time curves. Time-lapse movies were stabilized, maximum-projected (ImageJ), and terminal dendrites selected for analysis based on their separation from neighboring dendrites and the signal-to-background ratio. Terminal dendrites were selected throughout the arbor. Examples are shown in Figure 3 A. Extraneous objects were manually deleted.

To track the growing and shrinking tips, the algorithm determined the longitudinal centerline of the terminal dendrite and the location of its end for each frame. The central line was computed by fitting Gaussians to the cross-sectional intensity profiles at regular intervals along the backbone of the dendrite using:

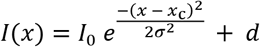

(Demchouk et al., 2011) where *x* is the position along a normal to the dendrite, *I*_0_ is the peak intensity value, *x_c_* is the center of the Gaussian, *σ* is the standard deviation and *d* is the measured camera offset (100) plus the background fluorescence. To compute the location of the tip, (*x*_tip_, *y*_tip_), we fit a 2-dimensional Gaussian function convolved with an error function using (Demchouk et al., 2011):

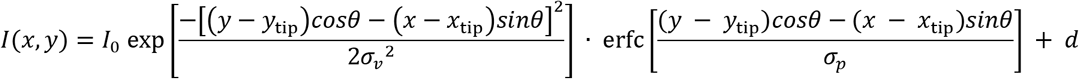

where *I*_0_ is the peak intensity, *θ* is the angular direction of the tip, and *σ_v_* and *σ_p_* are the standard deviations along the orthogonal and parallel directions. The length of the dendrite in each frame was determined by fitting the center line and the tip with a cubic spline. Length-time traces were smoothed with a median filter of size 3 to remove glitches.

To estimate the precision of our tracking algorithm, we used two different approaches. In the first, we tracked synthetic images of capped cylindrical tubes of known length and radius with fluorophores placed randomly on their surfaces (10% labeling density) and convolved with a point spread function (350 nm FWHM). To ensure that our algorithm could perform robustly under a wide range of signal-to-background ratios, we tested the tracking accuracy with decreasing signal to background ratios. The typical precision was ≪ 1 pixel (100 nm) even for low signal-to-background ratios (Ruhnow et al., 2011). (ii) We tracked the position of *in-vivo* dendritic tips that are in long-term paused states and found that the average standard deviation of length was ~0.1 μm (< 1 pixel, 0.1615 μm) as shown in Figure S2 D. This accuracy is comparable to and, in some cases, better than available software, such as, FIESTA (Ruhnow et al., 2011), JFilament (Smith et al., 2010), and Simple Neurite Tracer (Longair et al., 2011). Using a parallelized method, several hundred tips can be tracked simultaneously. A caveat of our method is that it can only track filaments that are reasonably free of extraneous objects, excessive noise, and have no breaks, discontinuities, or overlapping segments.

### Calculating velocities and transition rates

Fitting piecewise linear functions to the data is an ill-posed problem (Hansen and O’Leary, 1993) because a perfect fit can always be achieved with a large enough number of segments (equal to the number of data points minus one). To circumvent this problem, we defined a temporal resolution, *T* (in frames), that is necessary to distinguish a transition event from the noise in the data. Then, the maximum number of segments in each trajectory was calculated by dividing the total number of data points (total number of frames) by the temporal resolution. We used simulated data to estimate the temporal resolution that performs the best. We generated Markovian trajectories with known and realistic velocity distribution (±1.5 μm/min) and transition rates (0.5 min^-1^), similar to those shown the Figure 3. We then, added Gaussian white noise of standard deviation 0.25 μm to the trajectories to mimic the experimental noise. To analyze the trajectories, we used the following steps as shown in Figure S9:

i. We fit the trajectories with piecewise linear function considering *N*_data_/*T* as the initial number of segments (*N*_seg_).
ii. The velocity distribution (slope of the segments) was fitted to a lognormal-Gaussian-lognormal distribution:

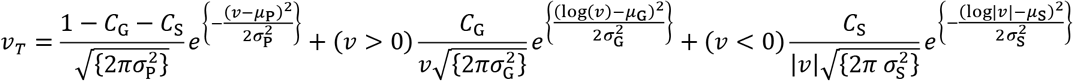

where the *C*’s are the normalization constants, the μ’s are the means, the σ’s are the standard deviations and the subscripts P, G, S stand for paused, growth, and shrinkage states respectively. The first term is a Gaussian and denotes the paused state, whereas 2^nd^ and 3^rd^ terms are log-normal distributions representing the growth and shrinkage states. After fitting the above velocity distribution, the intersections between the paused and growth distribution (*I*_G_) and the paused and shrinkage distribution (*I*_S_) were calculated, and the segments labeled using these two velocity thresholds.
iii. Consecutive segments with similar labels were then merged. This process decreased the number of segments (*N*^n^_seg_). The trajectories were then refitted with the new number of segments. Steps (ii) and (iii) were repeated until *N*^n^_seg_ stabilized.
iv. Finally, the transition rates were calculated by counting the total number of transitions from one state to another and then dividing that number by the total time spent in that state. For example, if *N*_GP_ is the total number of transitions from G to P state and *T*_G_ is the total time spent in the G state, then the transition rate from G to P is 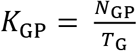.

Figure S9 shows the working protocol along with the validation of our analysis method. To estimate the optimal frame resolution, we plotted the root-mean-squared difference between the input and output transition rates as a function of the frame resolution. Our analysis clearly shows that temporal resolution of 6 frames generates the best results for rates ~0.5/min and noise ~0.25 μm. We used this frame resolution for all our tip dynamics data analysis.

### Average tip velocity & validation of segmentation

The segmentation yields a set of intervals, *i*, with associated distances, *d_i_*, durations, *t_i_* and velocities, *v_i_* = *d_i_*/*t_i_*. The distribution of velocities is fit to a lognormal-Gaussian-lognormal model *p*(*v*) with parameters *P*_G_, *μ*_G_, *σ*_G_, *P*_P_, 0, *σ*_P_, *P*_S_, *μ*_S_, *σ*_S_ such that

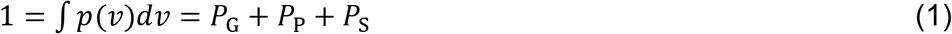

and

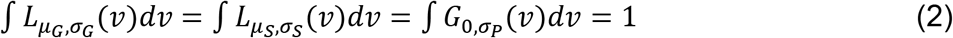

where *L_μ_G_,σ_G__*(*v*) and *L_μ_S_,σ_S__*(*v*) are the log-normal distributions corresponding to growth and shrinkage and *G*_0,*σ_P_*_(*v*) is the Gaussian distribution of the paused state. Transitions can only occur between unlike states. This imposes an important structure on the data: there are two threshold velocities *v_+_* and *v_* such that if the segment velocity *v_i_* > *v_+_* then it is assigned to be a growing segment. Likewise, if *v_i_* < *v_* it is a shrinking segment. The ones in the middle are paused.

The average velocity as:

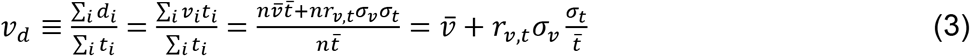

where 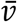 is the mean velocity, 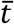t is the mean time, and *r_v,t_* is the Pearson correlation coefficient. We used the definition of the Pearson’s correlation coefficient *r_x,y_* = *σ_x,y_/σ_x_σ_y_* and cross-correlation 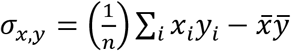 to calculate equation 3. The average velocity *v_d_* has two parts: 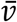, calculated assuming *t_i_* and *v_i_* are independent of each other, and a cross-correlation term, 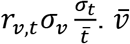 is given by:

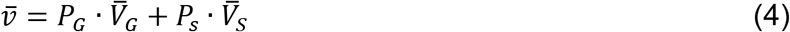

where 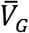 and 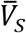 are the first moments of the lognormal velocity distributions for growth and paused states, and *P_G_* and *P_s_* can be calculated from the master equation associated with the transition matrix:

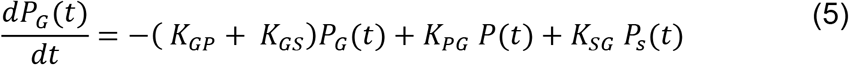

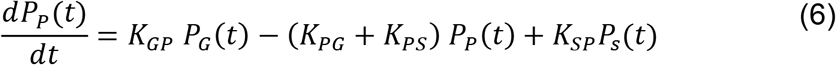

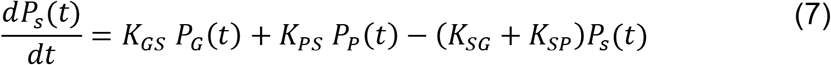

where *K_ij,j≠i_ i,j* = {*G,P,S*} are the transition rates. The steady state solution, assuming 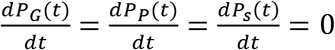 and using equation 1, is:

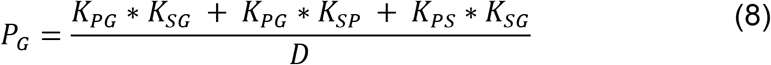

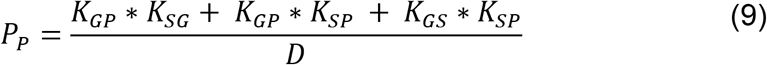

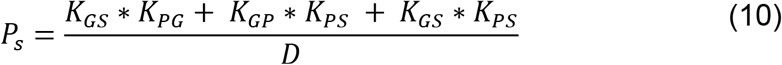

Where, 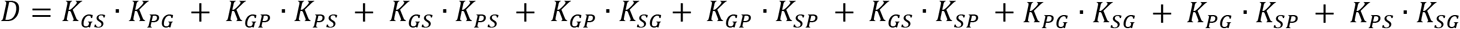.

Finally, the average velocity is:

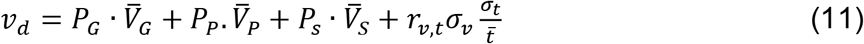

The average velocity is a key parameter, which controls the growth of the simulated arbor (Figure S5 D & E). The average velocity calculated in this way from the transition matrix agreed with that calculated directly from the raw traces at all developmental stages (Table 1) considering the Pearson correlation coefficient *r_v,t_* = 0 (Figure S3 L) This validates of our segmentation scheme.

### Computational model

The agent-based two-dimensional computational model of dendritic growth incorporated the fundamental processes that govern the morphogenesis of Class IV neurons: (i) branching, (ii) tip dynamics, and (iii) contact-based retraction (non-overlapping). We started our simulation with randomly oriented 2-4 branches emanating from the origin (cell body). Each branch is a filament, and points (*x,y*) are added at 0.1 μm (Δ*l*) intervals as the branch grows. The simulation is divided into 0.1-minute time steps (*Δt*). The details of individual processes are as follows:

#### Branching

Assuming the branching is a random process, we visit all the branches randomly and calculated the branching probability *P_b_* = 1 – *e^L_b_ω_b_Δ_t_^*, where *L_b_* is the length of the branch and *ω_b_* is the branching rate per unit time per unit length (Figure 4A). Then, a uniformly distributed unit random number Æ(0,1) is compared to *P_b_* to spawn a nascent branch from a random point on the mother branch. The branching angle is chosen randomly from the measured branch angle distribution which is distributed normally with a mean ~90° and standard deviation ~26° as shown in Figure 4B and Table S1. Each newly spawned branch is assumed to start in the growing phase with an initial length of 0.5 μm.

#### Tip Dynamics

Each branch with a free end (tip) follows a Markov process (after an initial lag, see below), transitioning between growing (G), paused (P), and shrinking (S) states with measured transition rates (*K_ij,j≠i_ i,j* = {*G,P,S*};) and velocities (*V*{*g,p,S*}) for free tips (Figure 3 D,E & Table 1). The transition dynamics is implemented using a standard ‘Monte-Carlo’ method. At each time step, the total probability of transition is calculated using *P_i_* = 1 – *e^-K_tot_Δt^*, where *K_tot_* is the sum of the transition rates from one particular state: *K_tot_* =. For example, the total transition rate from the growth state is *K_tot_ = K_GP_* + *K_GS_.* Subsequently, *P_i_* is compared with a uniform random number *R*(0,1) to implement the transition. If there is a transition, it happens maintaining the ratio *K_ij_/K_tot_*. After the transition, the tip is assigned with corresponding state velocity, the magnitude of which is randomly chosen from their respective velocity distributions (lognormal for growth and shrinkage and normal for paused) (Figure 3D). The growth process is implemented by adding new points to the existing branch tip at each time step, *Δt,* as follows:

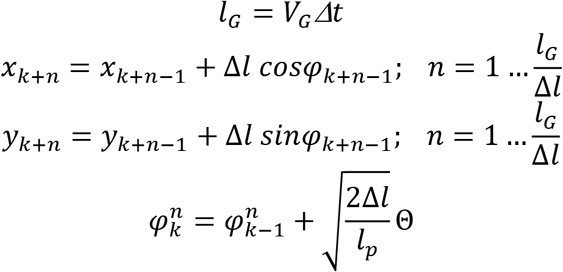

Where, *k* is the index of the last point in the previous time step and *l_p_* is the persistence length of the branches (set to 150 μm, Table S2). Additionally, the *Δl* value of the last point is adjusted if *l*_G_ is not an integer multiple of *Δl.* Θ is a unit Gaussian variable centered at zero.

While shrinking, points from the branches are removed until the shrinking length *l_s_ = v_s_Δt* is reached.

#### Contact-based retraction

It has been shown by several studies that Dscam molecules play an important role in the self-avoidance of dendritic tips in Class IV neurons (Matthews et al., 2007; Soba et al., 2007). To investigate the phenomenon, we measured the dynamics of the dendritic tips after a collision/contact event has occurred. Interestingly, we observed that the transition rates are altered after contact as shown in Table 1B and leads to an overall shrinkage of the dendritic tips. To implement this observation in the model, we assumed that the contact is achieved whenever a tip comes very close (<0.15 μm, ~average radius of branches, Table S2) to a nearby branch. Further, we used the altered tip dynamical parameters after a tip makes a contact (post-contact). Because it is difficult to measure how long the tips remain in their post-contact dynamics, we added this as a free parameter in the model (Table S2). It was chosen to be longer than the expected lifetimes of post-contact dendrites (10-15 minutes).

#### Boundary Condition

Individual Class IV neurons grow within the hemisegments (Grueber et al., 2002). We have experimentally measured the segment sizes in the AP and LR directions at different developmental stages as shown in Figure 1D. Linearly fits to the growing region (24 to 96 hr) defined the growth rates of the boundary used in our model. Because there is self-avoidance interaction between the neighboring neurons, we used a periodic boundary condition.

#### Initial condition

To avoid any ambiguity in the initial timepoint in the simulation, we divided the simulation process into two halves. We started our simulation at 14 hr with 2-4 randomly oriented branches of length ~15 μm and allowed them to grow with the 18 hr tip dynamics data (Table 1 A) and the 24 hr branching rate. When the dendritic tree reached the measured value of total branch number at 24 hr, we reassigned this time as 24 hr. In this way, we simulated larval morphogenesis.

#### Initial Lag

There is an unexpected paucity of dendrite deaths at short times. To analyze this, we calculated the survival probability of the branches in the following way:

Suppose dendritic tips are born and die throughout the observation period *T.* Divide *T* into small equal intervals *dt.* Let,

*f*(0) = Number{die in time [0, *dt*)}/Number{alive at time 0}
*f*(*dt*) = Number{die in time [*dt, 2*dt*)}/Number{alive at time *dt*}*
*f*(2*dt*) = Number{die in time [2*dt*, 3*dt*)}/Number{alive at time 2*dt*}.etc.

Then we can write the survival probability as:

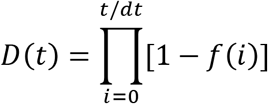

The survival probability of the experimentally observed dendritic tips was measured manually and then calculated by using the above formula. This is shown by the solid black line in Figure S4 A. The survival curve does not decay exponentially which led us to conclude that the tips have some initial period of sustained growth which we termed as initial lag *τ_lag_*. To estimate the amount of initial lag we simulated 1000 free tips with 48 hr tip dynamics data (because it is in the middle of the developmental time) and implemented an initial lag (*τ_lag_*) during which the tips did not switch into the paused or shrinking states (*K_GP_* = *K_GS_* = 0;*t* ≤ *τ_lag_*). We calculated the survival probability by dividing number of alive branches by the total number of branches. The survival probability increases with the initial lag *τ_lag_* as shown by the dotted lines in Figure S4A. The dark blue is the best fit to the real data (*τ_ļag_* = 0.3 min, Table S2) and we used this value in our model.

### Markovian tests of the tip trajectories

The tip dynamics is not a ‘Markovian process’. The non-Markovian traits are shown by the presence of initial lag during nascent branch formation (τ_lag_) and the change of dynamics after contact (Table S2). However, the dynamics is a first-order process as shown by the single exponential decay of phase durations as shown in Figure S3 A-I which points towards the fact that the dynamics don’t have any long -term memory. To confirm this, we calculated the state-state auto-correlation function (data not shown). The average autocorrelation function quickly becomes uncorrelated showing the absence of any long-term memory in the states.

## Acknowledgments

We thank all members of the Howard lab for many helpful discussions and suggestions. Special thanks to Dr. Ashley Arthur for proofreading the manuscript. We thank Hermann Cuntz, Damon Clark, and John Carlson for comments on an earlier version of the manuscript, and Chun Han and the Bloomington Drosophila Stock Center for fly stocks.

## Author contributions

SSh, SSu and JH designed the experiments based on early results by XL. SSh conducted the experiments and analyzed branching. SSu, OT, YT and JH conceptualized the models, which were implemented by SSu, who performed image and tip tracking analysis and wrote the source code for the *in-silico* model, based on earlier work by OT. SSh, SSu and JH wrote the manuscript.

## Data and materials availability

Data and code will be made publicly available.

## Supplementary Materials

### Supplementary Figures

**Figure S1:**
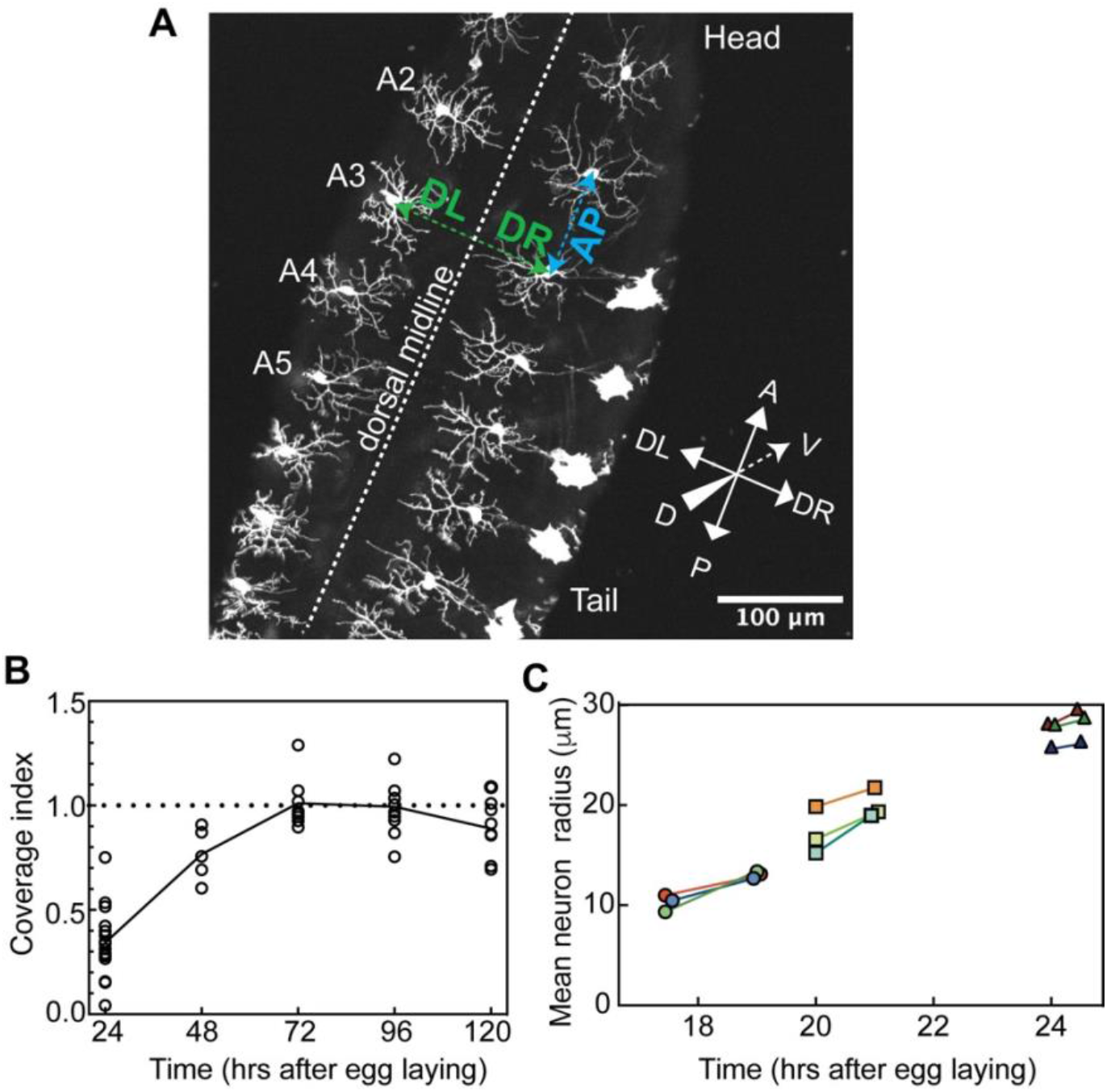
Definitions of the axes, the coverage index and neuronal growth controls. **A** Definitions of the anterior-posterior (AP) and dorsal left (DL) and dorsal right (DR) axes. This larva was imaged from the dorsal side up, which was adjacent to the coverslip surface closest to the objective. A2, A3, A4, and A5 correspond to the dorsal abdominal segments. The white dashed line is the dorsal midline. This larva is ~24 hr after egg lay (genotype - ;;ppkCD4-tdGFP). The AP length was measured as the distance between the cell bodies of the adjacent neurons on the anterior and posterior sides. The DL-DR length was measured as the distance between cell bodies in adjacent hemisegments (across the midline) corrected for the offset of the cell bodies, which are not in the centers of the cells but displaced away from the midline. For sake of simplicity we are calling DL-DR as LR **B** The coverage index over development time (*n*≥5 neurons). The coverage index is calculated as the ratio the dendrite area (AP cell width x LR cell width, from Figure 1D) divided by the dorsal hemisegment area (AP hemisegment width x LR hemisegment, width from Figure 1D). **C** Control showing that imaging does not perturb growth. The growth of cells was assessed by measuring the cell radius (calculated as √(area/π)) over time. Lines connect cells at the beginning and end of imaging. This shows that the imaging conditions do not retard growth.

**Figure S2:**
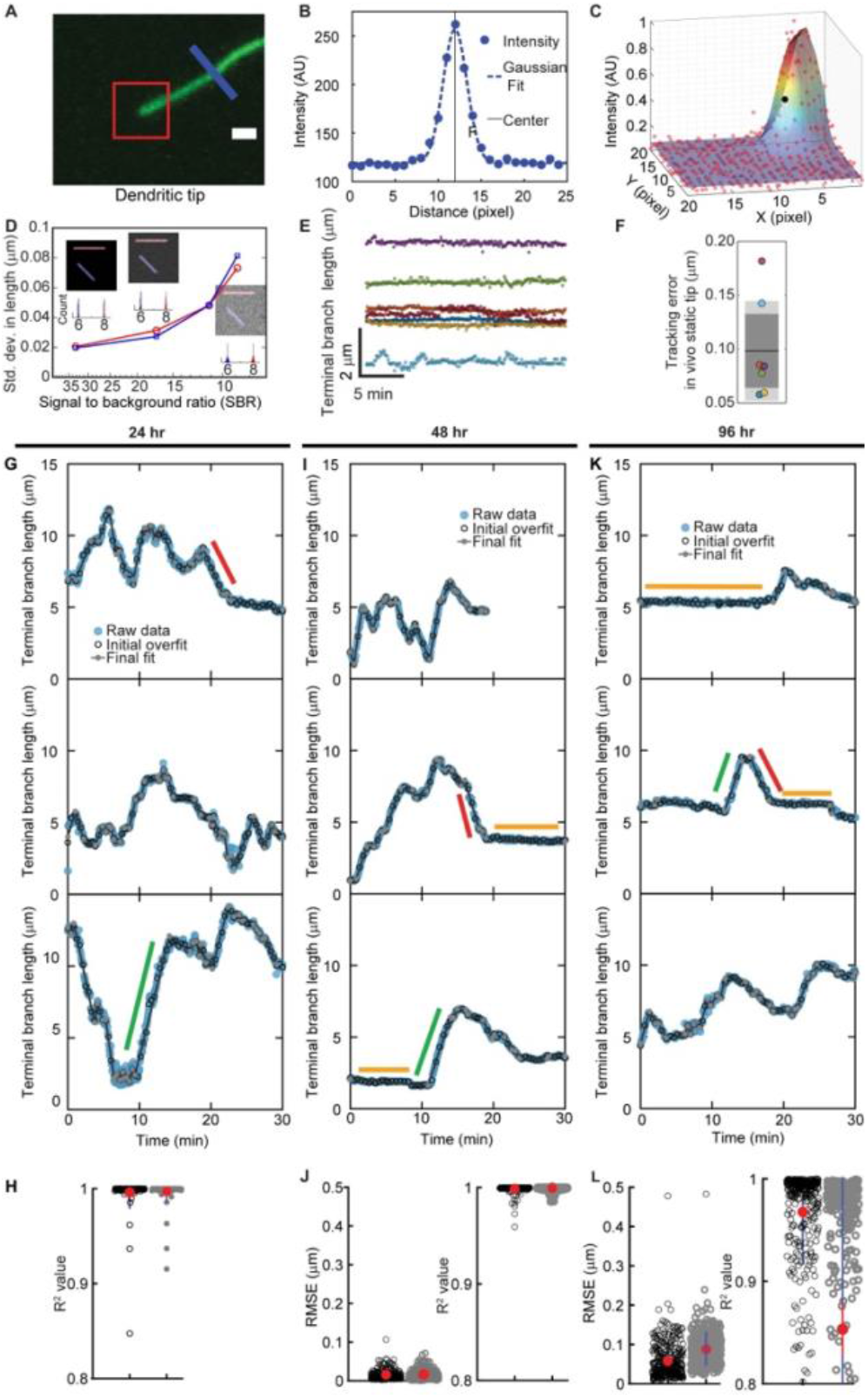
Tracking dendrite tips. **A** Example of a fluorescently labeled terminal dendrite. **B** The center of the dendrite was located by fitting the cross-sections (blue line in B) to a Gaussian. The precision is approximately 0.1 μm. **C** The position of the tip of the dendrite was calculated by fitting the end intensity profile (red box in A) to a 2D function corresponding to a Gaussian in the perpendicular direction and an error function the parallel direction (see Methods) **D** Montage of simulated images of cylindrical tubes (6 μm blue, 8 μm red) that are fluorescently labeled with 10% labeling density on the surface with signal-to-background ratio (SBR), defined as the mean signal divided by the standard deviation of background noise, varying from 33 (left) to 9 (right). The pixel size is 100 nm. The measured length distribution is shown in the bottom panel (200 independently generated images for each SBR). **E** The lengths of several live-imaged dendrites that were in their paused state as a function of time. **F** The standard deviation of the measured lengths in E is shown. The accuracy is high even for live imaging condition. **G** The standard deviation of the tracked lengths is ~1 pixel (108 nm). Examples of tracked dendritic length as a function of time: **G** 24 hr. **I** 48 hr. and **K** 96 hr. The green, orange, and red lines denote examples of growing, paused, and shrinking states. The tips tend to spend more time in the paused state over developmental time. **H**, **J**, and **L** show the statistics of the piecewise-linear fitting.

**Figure S3:**
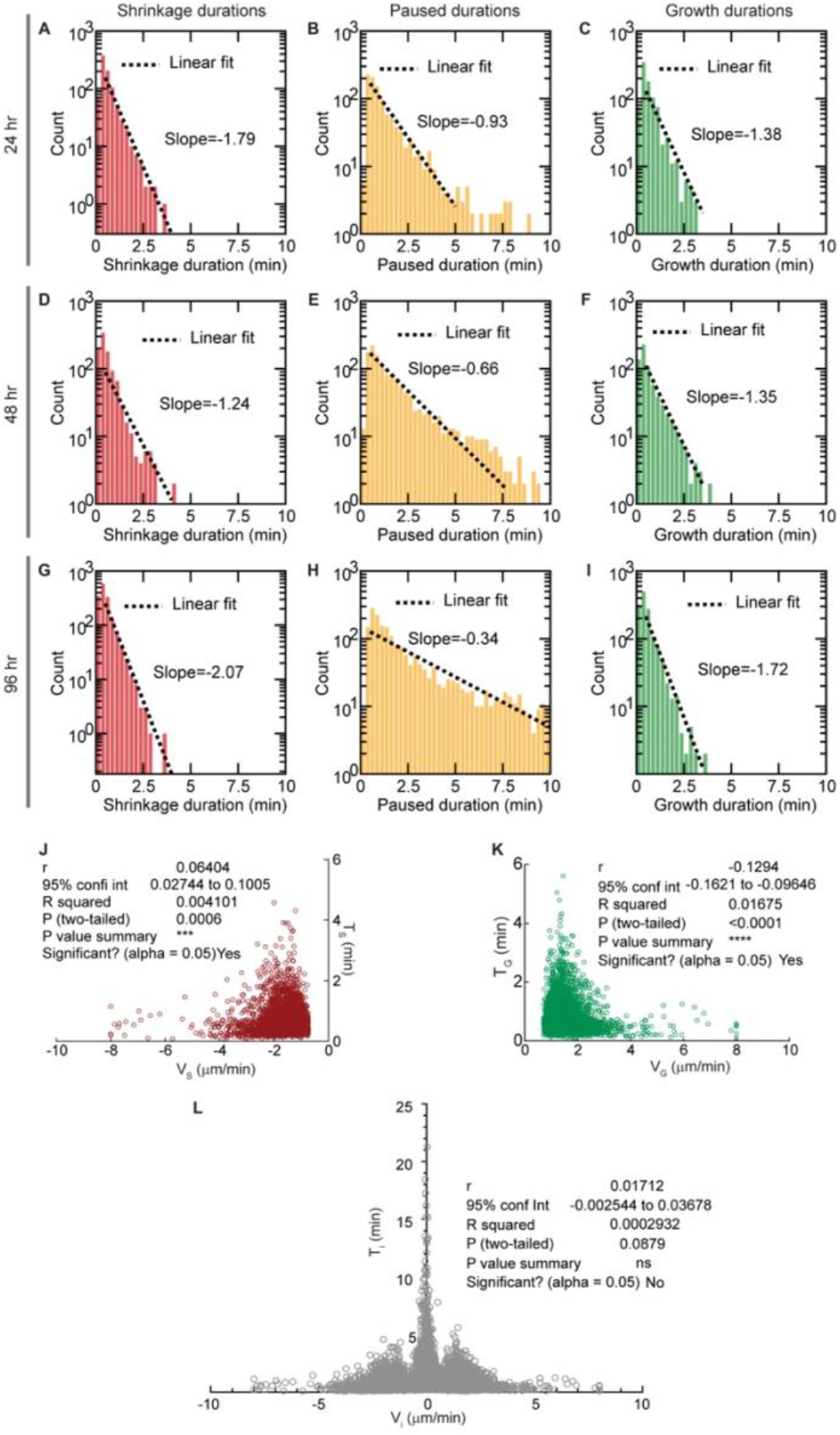
Correlation between state velocities and lifetimes and state durations. The distribution durations for the shrinking (S), paused (P), and growing (G) states at 24 hr (**A-C**), 48 hr (**D-F**), and 96 hr (**G-I**) plotted using semi-log axes. The distributions are very close to exponentials (dotted lines) expected if switching among the states is first order. The slope of the dotted lines is the inverse of the lifetimes spent in the states: for example, at 24 hr, the sum of the transition rates from growing state is (*K*_GP_ + *K*_GS_ = 0.696 + 0.509 = 1.205 min^-1^ (Table 1 A), close to the slope of the lifetime distribution of 1.38 min^-1^ (F). **J** The correlation between shrinking velocities (*V*_S_) and shrinking lifetimes (*T*_S_) shows a significant correlation with Pearson’s correlation coefficient r=0.064. **K** Similarly, a significant correlation is observed between growth velocities (*V*_G_) and growth lifetimes (*T*_G_). **L** Pearson’s correlation coefficient (*r*) between state velocities (*V_i_*) and lifetimes (*T_i_*)) for 18-20 hr data. The value of *r* is small (0.017) and there is no significant correlation between *V_i_* and *T_i_*.

**Figure S4:**
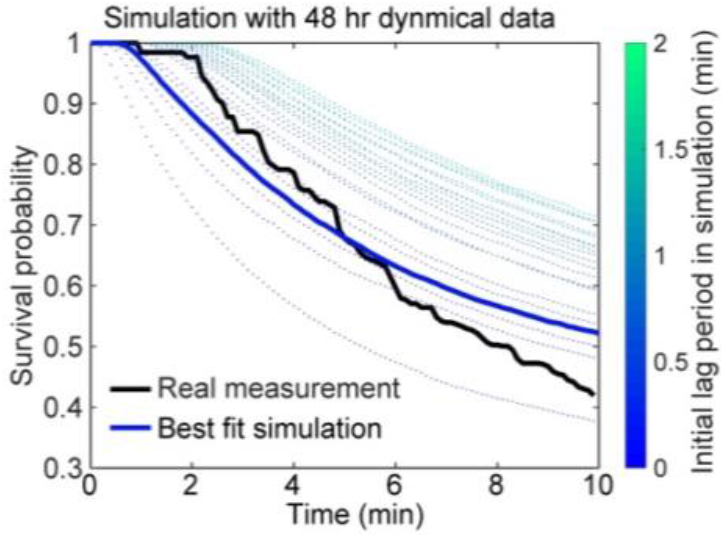
Evidence for persistent growth after birth. We manually measured the time between branch initiation and branch death at different stages of the larva (24, 48,72, and 96 hr AEL, 3 movies for each stage). From these data, we calculated the survival probability by dividing the number of alive branches by the total number of branches using the formula described in the Methods. The black line is the average survival probability of the real dendrite tips. The survival probability does not start decaying exponentially as one might expect if it were a Poisson process. Rather, it shows some initial lag. This observation led us to believe that branch initiation is not a simple Poisson process. To estimate the initial lag period, we simulated 1000 branches with initial length 0.5 μm and implemented a lag time (*τ_lag_*) by preventing the tips to switch into the paused or shrinkage state (*K*_GP_ = *K*_GS_ = 0;t ≤ *τ_lag_*). A branch is deleted in the simulation when its length is <0.1 μm. The survival probability increases with the initial lag *τ* as shown by the dotted lines. The dark blue is the best fit to the real data (*τ_lag_* = 0.3 min).

**Figure S5:**
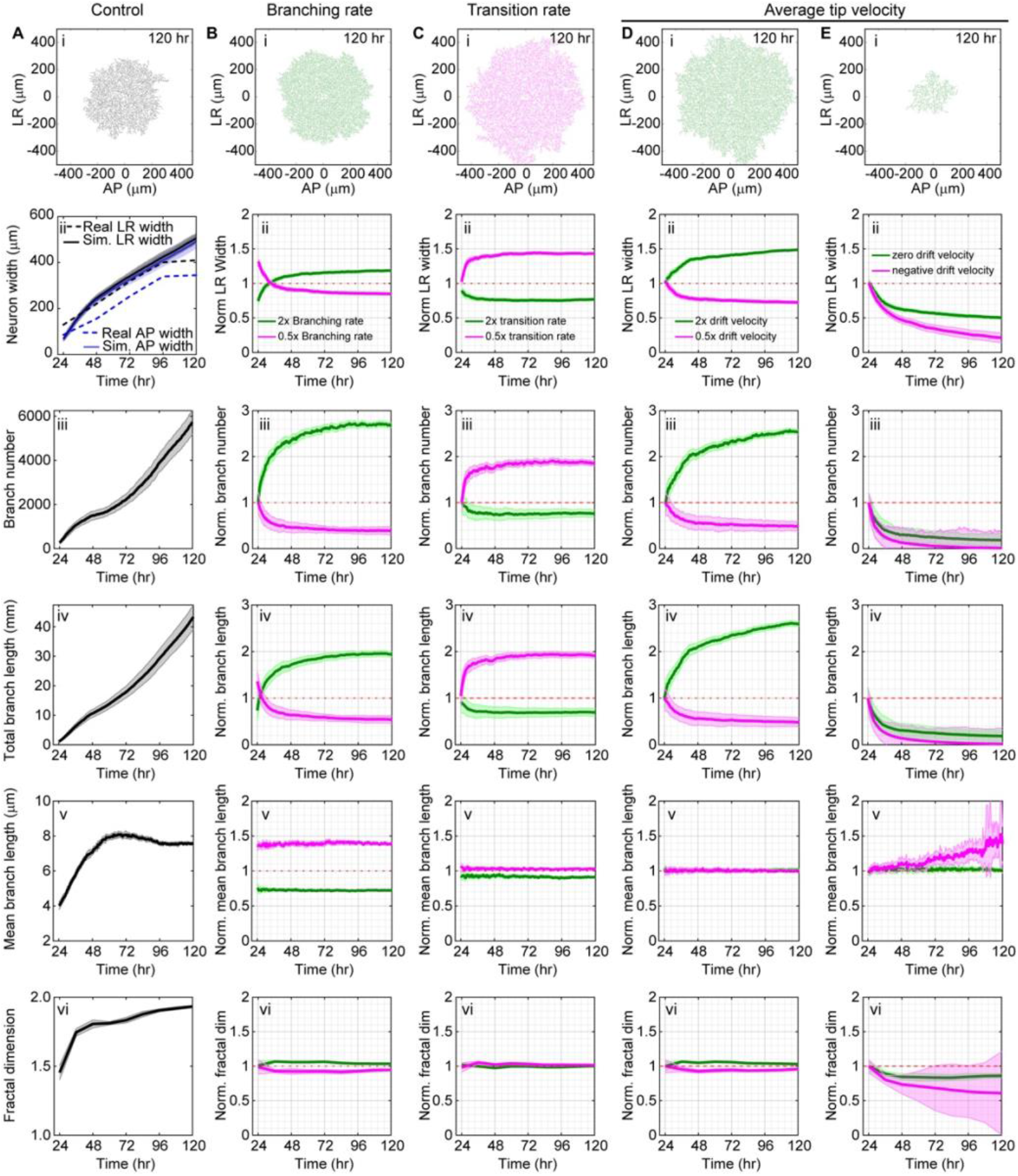
Sensitivity of morphology to branching and growth parameters. **A** Control with parameters from Table 1 and Tables S1-2 without any boundary restriction. The black and blue dashed lines represent LR and AP widths respectively. The solid lines represent the simulated segment sizes over development. The simulation shows that initially (24-48 hr) the neurons grow faster than the real segment and then grow with a constant rate equal to the segment growth rate (~0.06 μm/min) until 96 hr. The segment widths saturate after 96 hr even without a boundary. **B** Branching rate was doubled (green) and halved (magenta) compared to the control, keeping all other parameters unchanged. All arbor properties are normalized by the respective unconstrained controls. Fold change is plotted against time for (ii) arbor size, (iii) branch number, (iv) branch length, (v) mean branch length, and (vi) fractal dimension. **C** All transition rates were doubled (green) and halved (magenta): this leads to a decrease and increase in the variability of growth. **D** The mean tip velocity (drift) was increased (green) and decreases (magenta) 2-fold. **E** The average (drift) velocity was set to zero (green) and a negative value (−0.02 /min, magenta). Shaded regions represent standard error of mean.

**Figure S6:**
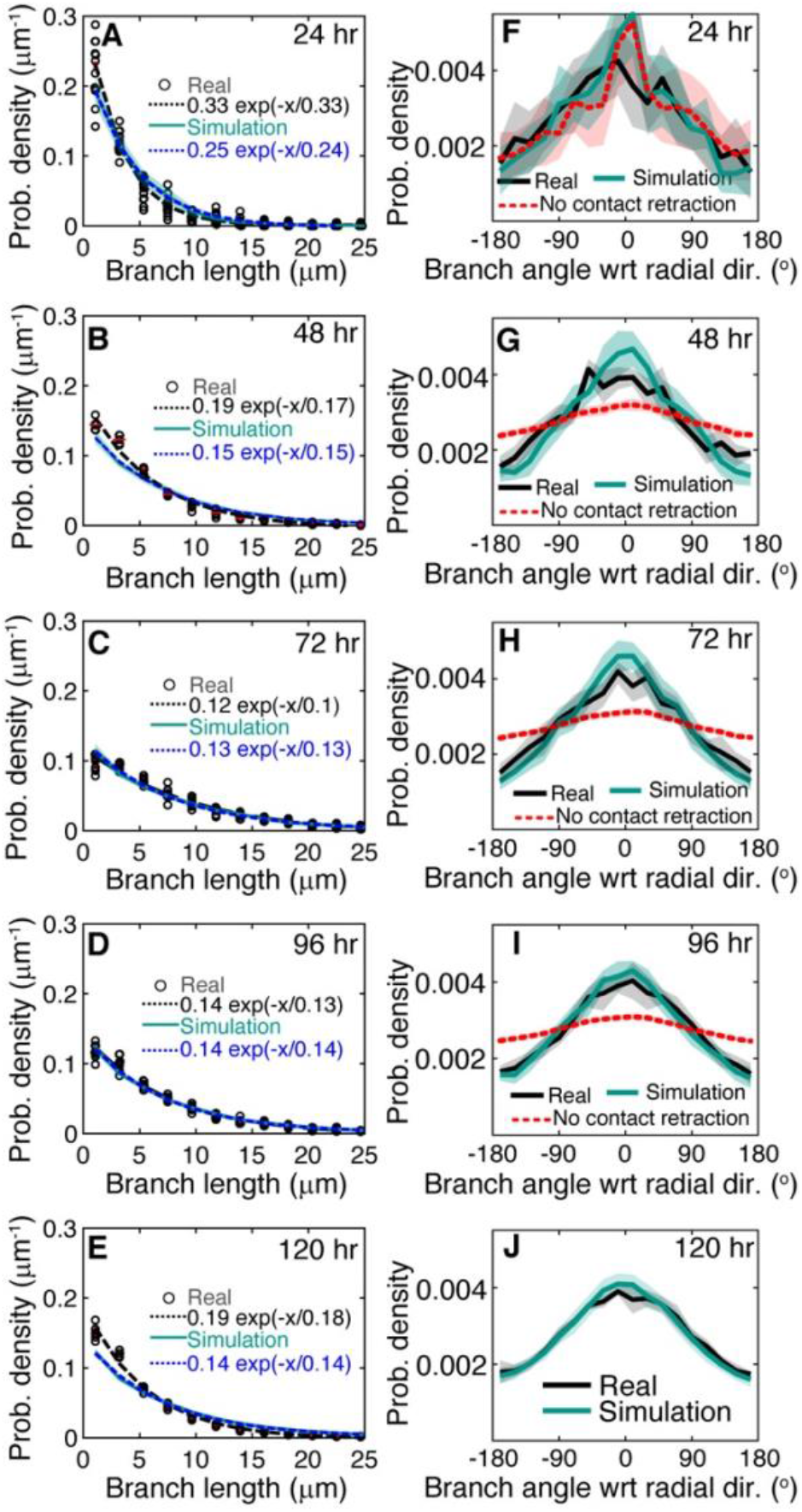
Branch length and radial orientation distributions. **A-E** Branch length distribution over different developmental stages for real and simulated arbors with exponential fits (dashed lines). **F-J** Radial orientation of branches over developmental time for real and simulated arbors. The branches are preferentially oriented in the radial direction. This preference is due to contact-based retraction. The dotted curves show diminished radial preference when branches are paused after contact in the simulation. Shades represents standard deviations.

**Figure S7:**
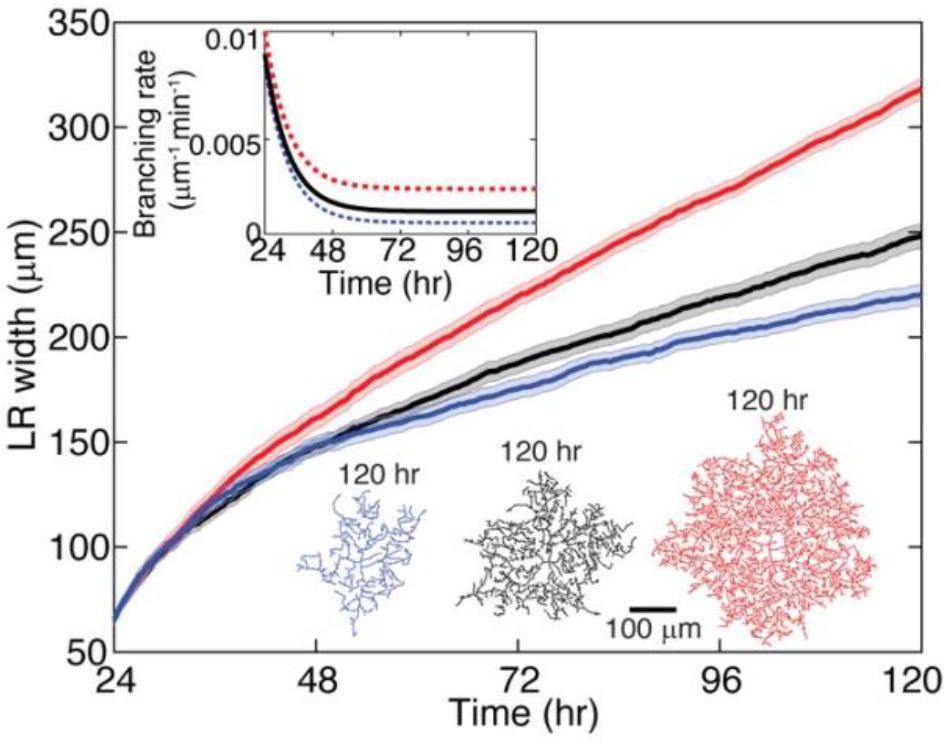
Branching drives arbor expansion when the net tip growth is zero. To understand the relationship between the short-term dynamics (the growth-shrink-pause dynamics including branching) and the long-term formation of stable branches, we explored our simulation keeping the net growth of tips at zero (meaning there is no net growth from G-P-S dynamics) and varied the branching rate (as shown by the red, black and blue lines in the top inset). We observed, even for zero net growth, that the dendritic arbor grows in size as shown by the LR widths (different colored arbors correspond to the differently colored branching rates in the top inset). The bottom panel shows the color-coded final arbor sizes.

**Figure S8:**
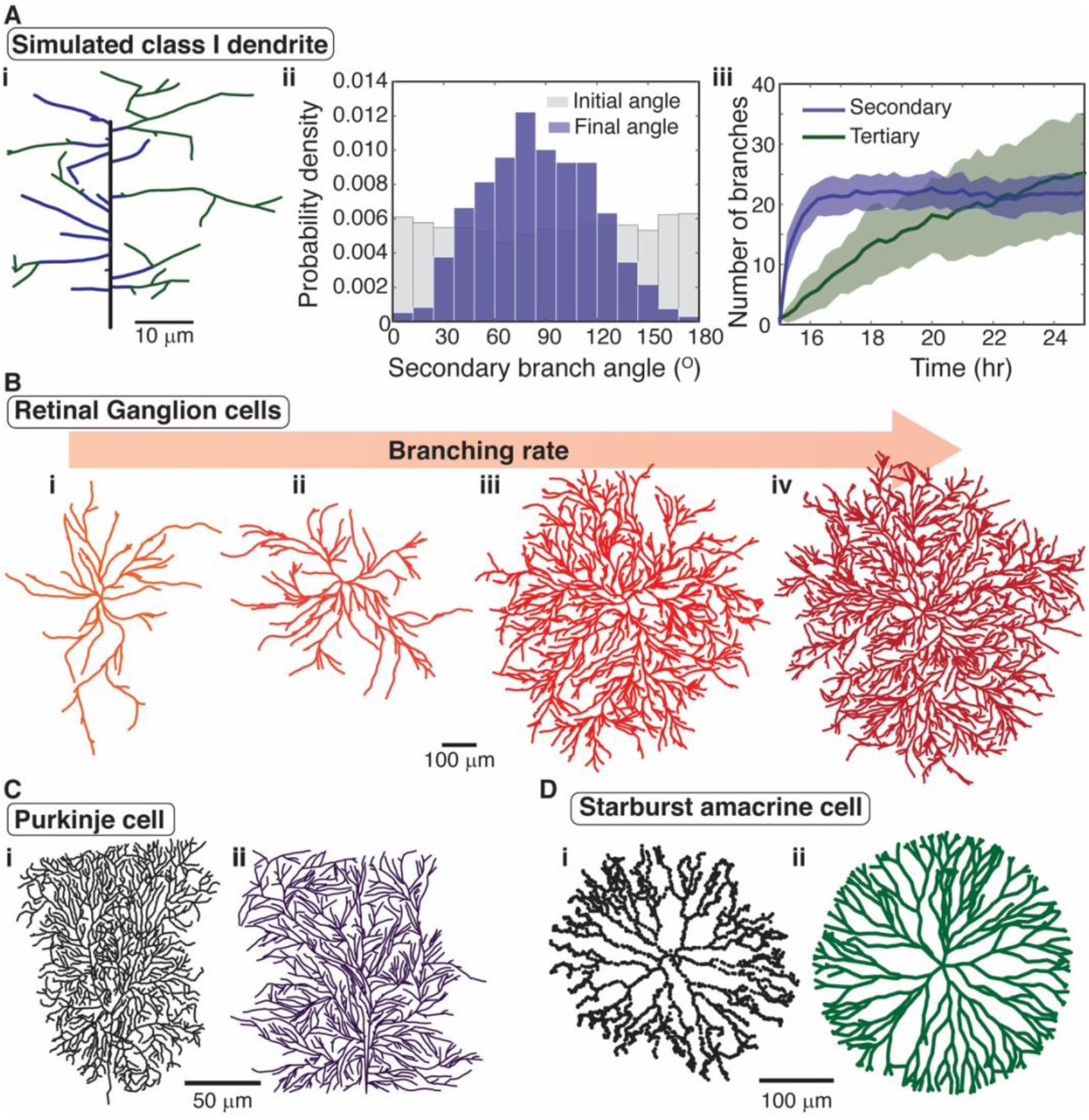
Generalization of our model to other systems. **A** (i) A representative simulated *Drosophila* class-I dendrite at 25 hr. Class-I dendrites were simulated by initializing the model with a single static primary branch and then allowing branching from primary and secondary branches with rates 0.05exp(-*t*/5) + 0.005 μm^-1^min^-1^ and 0.005exp(-*t*/5) + 0.0005 μm^-1^min^-1^) respectively, where *t* is time in hrs. (ii) The simulation recapitulates one of the key findings in (Palavalli et al., 2021) namely that the secondary branches are orthogonal to the primary branch (blue histogram peaking around 90 °) even though the initial angles were uniformly distributed (gray). This is a consequence of contact-based retraction. (iii) The number of secondary and non-secondary branches approaches 22 and 30 at long times respectively, in accordance with data from (Palavalli et al., 2021). **B** Different retinal ganglion cells were simulated using different branching rates and a small branching angle (45° relative to the direction of the mother). The morphologies are similar to those of marmoset retinal ganglion cells (Masri et al., 2019). **C** A real Purkinje cell (i) ((Murru et al., 2019), raw data downloaded from NeuroMorpho.org) was simulated using slow growth of dendritic tips and complete retraction after contact to recapitulate the locally parallel branch orientations (ii). **D** An example of a real starburst amacrine cell (i) ((Bloomfield and Miller, 1986), raw data downloaded from NeuroMorpho.org) and a simulated cell (ii) in which it was necessary to replace lateral branching with tip bifurcation to recapitulate the observed morphology.

**Figure S9:**
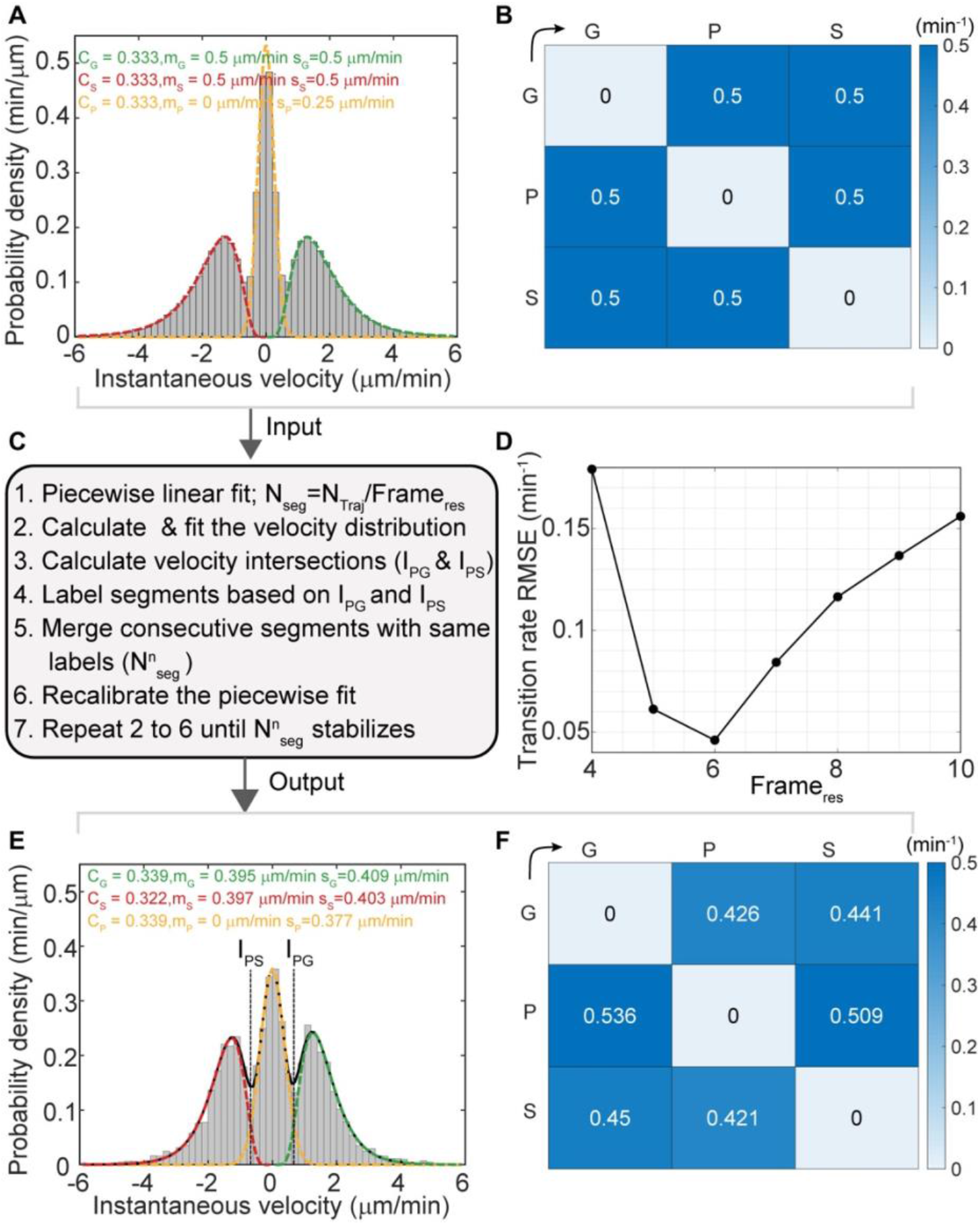
Validation of trajectory analysis method. We simulated 200 Markov trajectories with realistic input parameters shown in **A** and **B** and then added Gaussian white noise on the individual points of the trajectories. We used the transition rate as 0.5 /min because of the observed fact that the individual states last ~1 minute. **C** The flowchart of the trajectory analysis method. **D** We varied the Frame resolution to find the optimal resolution. The root-mean-squared error between the input and output transition rate matrix is plotted as a function of frame resolution. Frame resolution of 6 provides the best result and we chose this value for all our analyses. **E & F** The output velocity distribution and transition rate matrix using frame resolution 6. Our method of analysis produced a faithful reproduction of the input parameters.

### Supplementary Tables

**Table S1:** Branching rates.

| Age (hr) | Total branching rate<sup>a</sup> (min<sup>-1</sup>) (Mean ± SD) | Linear branching rate<sup>b</sup> (μm<sup>-1</sup>·min<sup>-1</sup>) (Mean ± SD) | Branching angle<sup>c</sup> (°) (Mean ± SD) | Numbers (rates, angles) |
| --- | --- | --- | --- | --- |
| 18 (E) | 4.26 ± 0.59 | 0.0109 ± 0.0015 | 85 ± 25 | *n* = 9, 5 |
| 24 (L1) | 7.59 ± 1.52 | 0.0095 ± 0.0017 | 88 ± 26 | *n* = 9, 7 |
| 36 (L1) | 5.77 ± 2.67 | 0.0031 ± 0.0009 | 85 ± 25 | *n* = 6, 7 |
| 48 (L2) | 4.86 ± 1.59 | 0.0019 ± 0.0007 | 86 ± 26 | *n* = 6, 4 |
| 72 (L2) | 8.31 ± 2.37 | 0.0011 ± 0.0004 | 90 ± 25 | *n* = 6, 5 |
| 96 (L3) | 11.12 ± 2.52 | 0.0011 ± 0.0006 | 91 ± 25 | *n* = 6, 4 |

<sup>a</sup>Over the entire dendrite arbor
<sup>b</sup>Per total dendrite length
<sup>c</sup>Angle between dendrites is zero in the distal direction of the mother.
Standard deviation (SD).

**Table S2:**
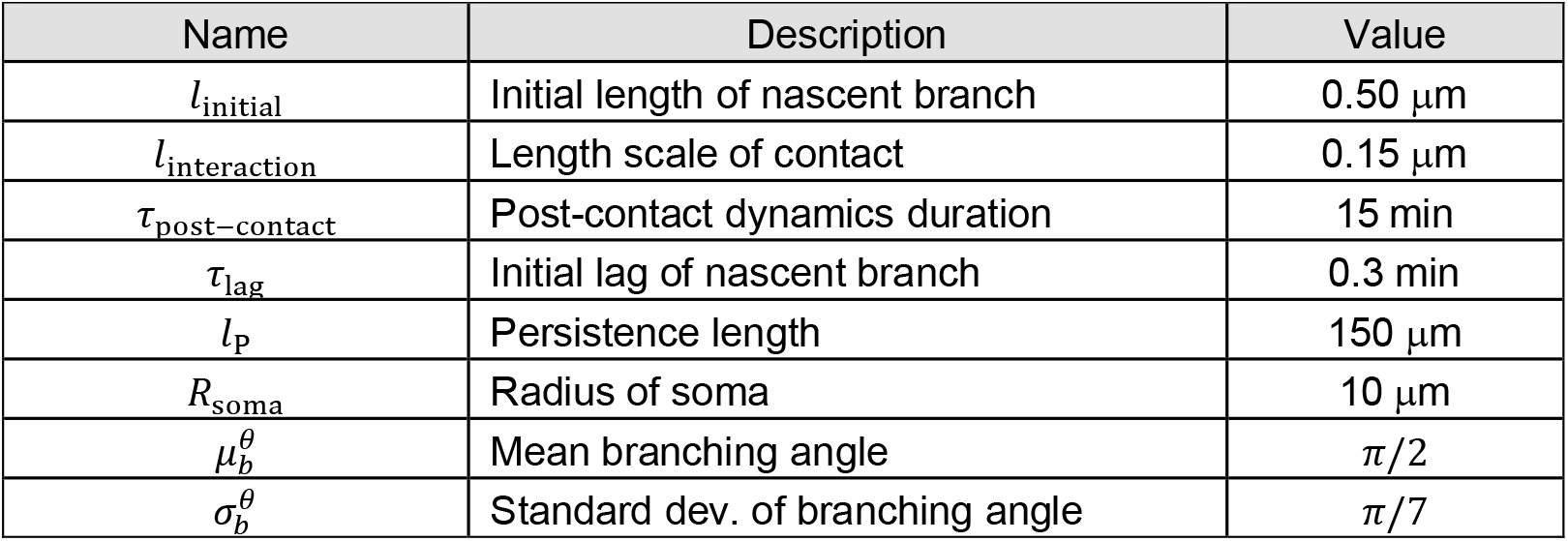
Model parameters.

**Table S3:**
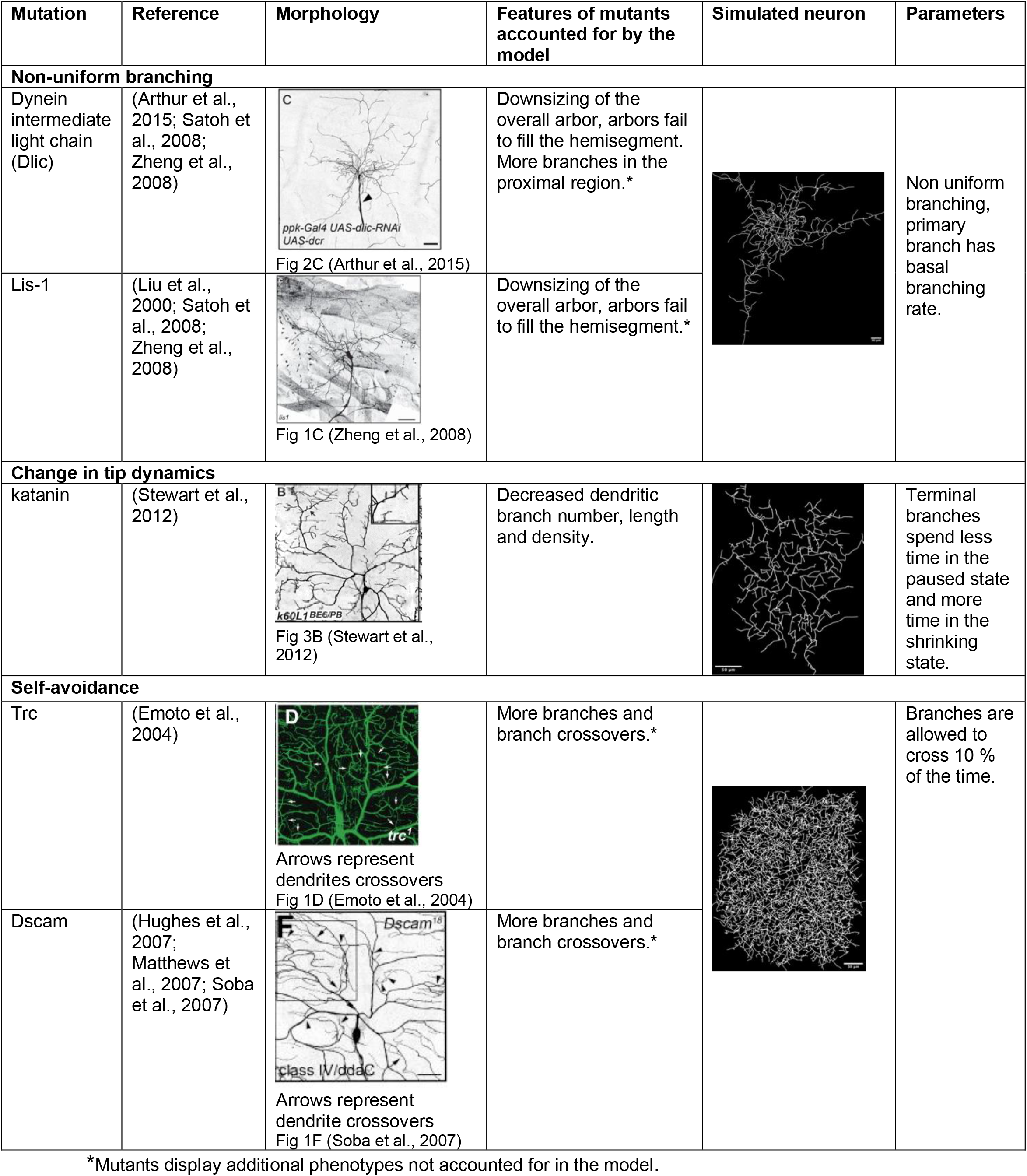
Mutations and morphologies.

| Mutation | Reference | Morphology | Features of mutants accounted for by the model | Simulated neuron | Parameters |
| --- | --- | --- | --- | --- | --- |
| **Non-uniform branching** |  |  |  |  |  |
| Dynein intermediate light chain (Dlic) | (Arthur et al., 2015; Satoh et al., 2008; Zheng et al., 2008) | Fig 2C (Arthur et al., 2015) | Downsizing of the overall arbor, arbors fail to fill the hemisegment. More branches in the proximal region.* |  | Non uniform branching, primary branch has basal branching rate. |
| Lis-1 | (Liu et al., 2000; Satoh et al., 2008; Zheng et al., 2008) | Fig 1C (Zheng et al., 2008) | Downsizing of the overall arbor, arbors fail to fill the hemisegment.* |  |  |
| **Change in tip dynamics** |  |  |  |  |  |
| katanin | (Stewart et al., 2012) | Fig 3B (Stewart et al., 2012) | Decreased dendritic branch number, length and density. |  | Terminal branches spend less time in the paused state and more time in the shrinking state. |
| **Self-avoidance** |  |  |  |  |  |
| Trc | (Emoto et al., 2004) | Arrows represent dendrites crossovers<br>Fig 1D (Emoto et al., 2004) | More branches and branch crossovers.* |  | Branches are allowed to cross 10 % of the time. |
| Dscam | (Hughes et al., 2007; Matthews et al., 2007; Soba et al., 2007) | Arrows represent dendrite crossovers<br>Fig 1F (Soba et al., 2007) | More branches and branch crossovers.* |  |  |

\*Mutants display additional phenotypes not accounted for in the model.

### Supplementary movies

**Movie S1.**
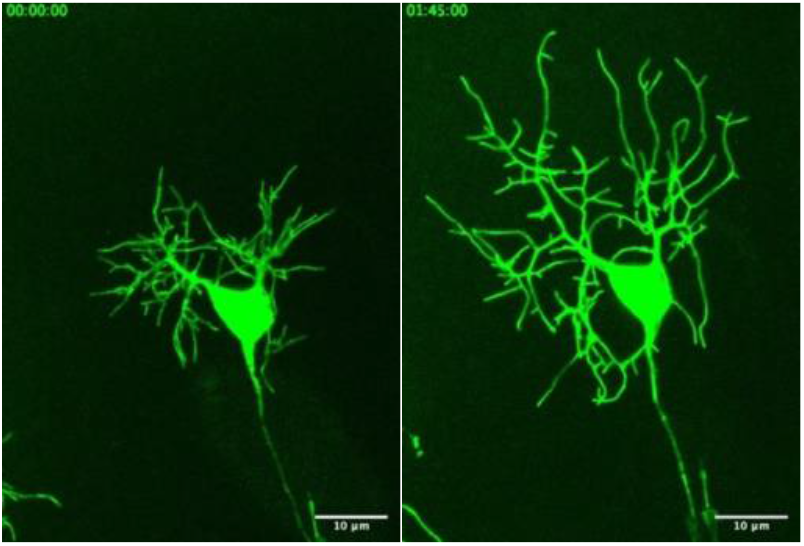
Time-lapse movie of a growing neuron: Time lapse movie of an fast embryonic neuronal growth at 17.5hr AEL was acquired using a spinning disk confocal microscope. The movie was full-frame (2048x 2048 pixels) and a complete stack of images (7um) was produced every 5 mins interval. Genotype of embryo was *;;ppkCD4-tdGFP*.

**Movie S2.**
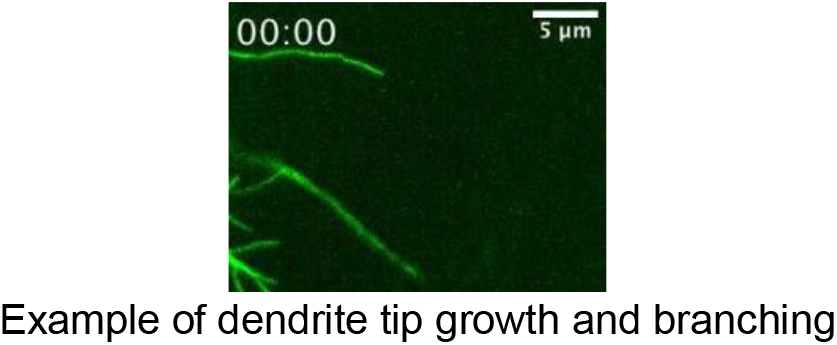
Tip growth and branching: Time lapse movie at 24hr AEL was acquired using a spinning disk confocal microscope. A cropped stack of images (7um) was produced every 5 seconds interval. Genotype of larvae was *;;ppkCD4-tdGFP*.

**Movie S3.**
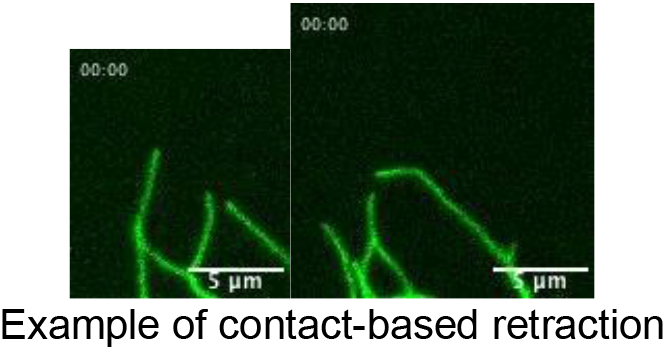
Self-avoidance and shrinkage: Time lapse movie at 24hr AEL were acquired using a spinning disk confocal microscope. A cropped stack of images (7um) was produced every 5 seconds interval. Genotype of larvae was *;;ppkCD4-tdGFP*.

**Movie S4.**
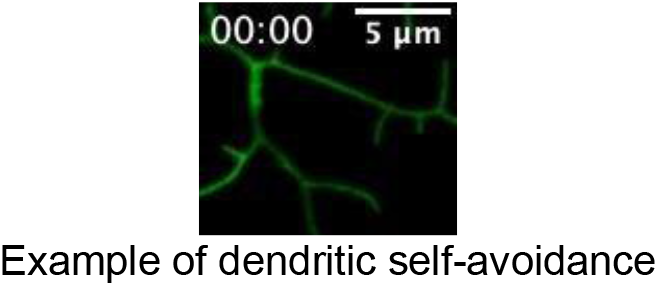
Self-avoidance and growth: Time lapse movie at 24hr AEL was acquired using a spinning disk confocal microscope. A cropped stack of images (7um) was produced every 5 seconds interval. Genotype of larvae was *;;ppkCD4-tdGFP*.

**Movie S5.**
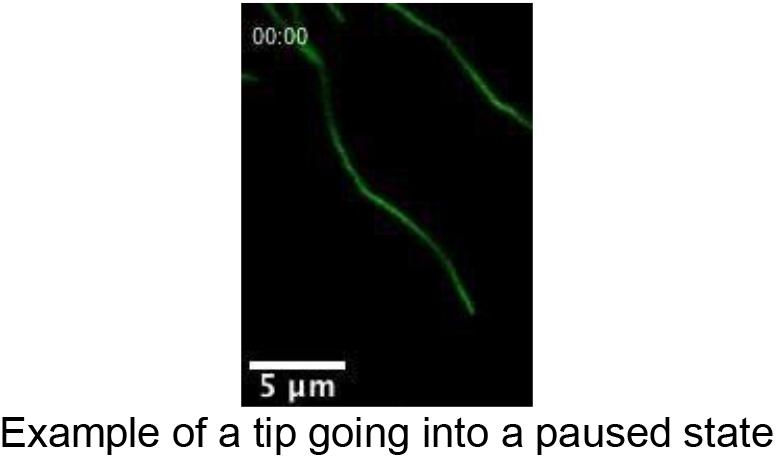
Tip pause: Time lapse movie at 24hr AEL was acquired using a spinning disk confocal microscope. A cropped stack of images (7um) was produced every 5 seconds interval. Genotype of larvae was *;;ppkCD4-tdGFP*.

**Movie S6-10.**
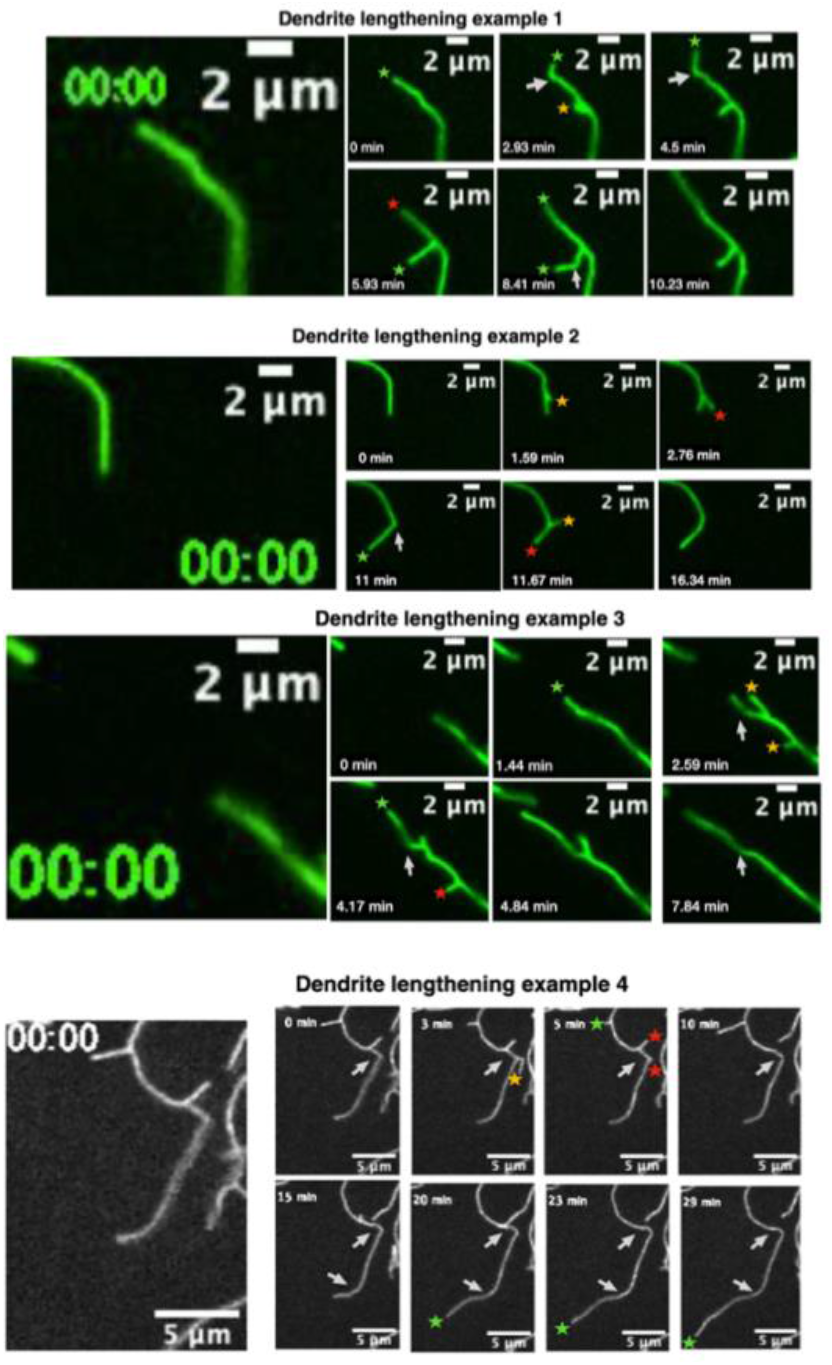
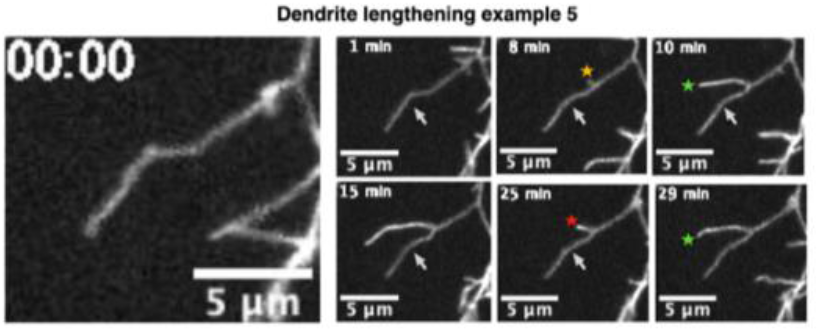
Tip growth, bending, and branching. Dendrite lengthening is likely due to the addition of materials at the dendrite tip. The green stars show dendrite tip lengthening, the yellow star is birth of new branch, and the red star is a shrinkage event. White arrows point to the bending of growing tips. In example two, where the branch disappears, a sharp bend smoothens over time. In examples 3,4, and 5, the white arrows correspond to structural features such as branches and bends that remain fixed during growth and shortening. All time lapse movies shown above were acquired from different 24hr larvae using spinning disk confocal. Genotype of all larvae was *;;ppkCD4-tdGFP*.

